# Accelerated social representational drift in the nucleus accumbens in a model of autism

**DOI:** 10.1101/2023.08.05.552133

**Authors:** Pingping Zhao, Xing Chen, Arash Bellafard, Avaneesh Murugesan, Jonathan Quan, Daniel Aharoni, Peyman Golshani

**Affiliations:** Department of Neurology, David Geffen School of Medicine, University of California; Los Angeles, Los Angeles, CA, USA; West Los Angeles Veteran Affairs Medical Center; Los Angeles, CA, USA; Intellectual and Developmental Disabilities Research Center, University of California; Los Angeles, Los Angeles, CA, USA

## Abstract

Impaired social interaction is one of the core deficits of autism spectrum disorder (ASD) and may result from social interactions being less rewarding. How the nucleus accumbens (NAc), as a key hub of reward circuitry, encodes social interaction and whether these representations are altered in ASD remain poorly understood. We identified NAc ensembles encoding social interactions by calcium imaging using miniaturized microscopy. NAc population activity, specifically D1 receptor-expressing medium spiny neurons (D1-MSNs) activity, predicted social interaction epochs. Despite a high turnover of NAc neurons modulated by social interaction, we found a stable population code for social interaction in NAc which was dramatically degraded in Cntnap2^-/-^ mouse model of ASD. Surprisingly, non-specific optogenetic inhibition of NAc core neurons increased social interaction time and significantly improved sociability in Cntnap2^-/-^ mice. Inhibition of D1- or D2-MSNs showed reciprocal effects, with D1 inhibition decreasing social interaction and D2 inhibition increasing interaction. Therefore, social interactions are preferentially, specifically and dynamically encoded by NAc neurons and social representations are degraded in this autism model.

## Introduction

To survive and thrive, animals need to sense, process and interpret incoming social cues and respond with appropriate behaviors in a context-dependent manner (*1*). Yet, how neural activity represents social information to drive successful decision-making and whether these representations are stable are still poorly understood. The social motivation theory of autism posits that social interactions are inherently rewarding and intrinsically motivated and that individuals with autism find social interactions less rewarding or motivating. Therefore, an individual with autism is less likely to orient toward, seek, or maintain social interactions because of changes in social motivational circuits (*2*, *3*). The nucleus accumbens (NAc), a key node in the mesocorticolimbic pathway, is critical for reward processing and motivated behavior (*4*, *5*). Whether and how distinct neuronal populations within NAc reliably encode social interaction and whether these neuronal populations are differentially activated in autism is unclear.

The NAc mainly consists of D1 receptor-expressing medium spiny neurons (D1-MSNs) and D2 receptor-expressing neurons (D2-MSNs) (*4*, *5*), with each cell type playing a distinct role in a range of reward behaviors (*6–9*). How these specific cell types encode and differentially motivate social interactions is still controversial (*10*, *11*). As social interactions evolve over time and previous interactions impact future ones, it is critical to examine activity not only within a single epoch but across sessions and days. Therefore, how long-term neural representation of social interaction in NAc at both the single cell and population levels are maintained is a fundamental question for understanding related circuits. Disturbance to the long-term stability of social representations may be a fundamental mechanism driving deficits in social interactions in ASD.

## Results

### NAc neurons are preferentially modulated by social interaction

Social interaction is rewarding for many species (*11–13*). To understand how social interactions are represented in the NAc, the main hub for reward circuitry, we performed *in vivo* calcium imaging using miniaturized microscopy to monitor neural activity in the NAc core during alternate sessions of social and object interactions in an open arena. Imaging was performed in adult male mice (11-12 weeks of age) as they interacted with other male mice that were 7-9 weeks of age (to avoid intruder aggression). Social interaction and object exploration were performed in a counterbalanced manner. Calcium signals and animal behavior were recorded simultaneously (Fig. 1A). We observed significantly more bouts of social interaction than object exploration, and mice spent a greater proportion of time interacting with the social target than exploring the object (Fig. 1B). Using miniscope calcium imaging, we recorded neural activity of individual NAc neurons during free behavior (Fig. 1C, D). Using receiver operating characteristic (ROC) analysis (*14*, *15*), recorded neurons were identified as significantly excited cells (social excited (SE) or object excited (OE) cells), significantly inhibited cells (social inhibited (SI) or object inhibited (OI) cells) or non-modulated cells (social non-modulated (SN) or object non-modulated (ON)) (Fig. 1G, H). Overall, we identified significantly more neurons modulated by social interaction (SE cells and SI cells) than those modulated by object interactions (OE cells and OI cells) (20.38±3.01% vs 11.48±1.77%, mean±SEM) (Fig. 1K). Specifically, over 80% of social modulated neurons were social excited cells (SE: 16.67±2.80%; SI: 3.72±1.27% of total cells, mean±SEM). The proportion of social excited neurons was substantially larger than that of social inhibited cells, object excited cells, and object inhibited cells (16.67±2.80% vs 3.72±1.27% / 5.35±1.70% / 6.13±1.60%, mean±SEM) (Fig. 1L). The overlap between social cells and object cells was very low (2.7%) in NAc (Fig. S1A). Overall, these results show that NAc neurons are preferentially modulated by social interactions, with mutually exclusive subpopulations that are engaged or inhibited by social stimuli.

**Fig. 1.**
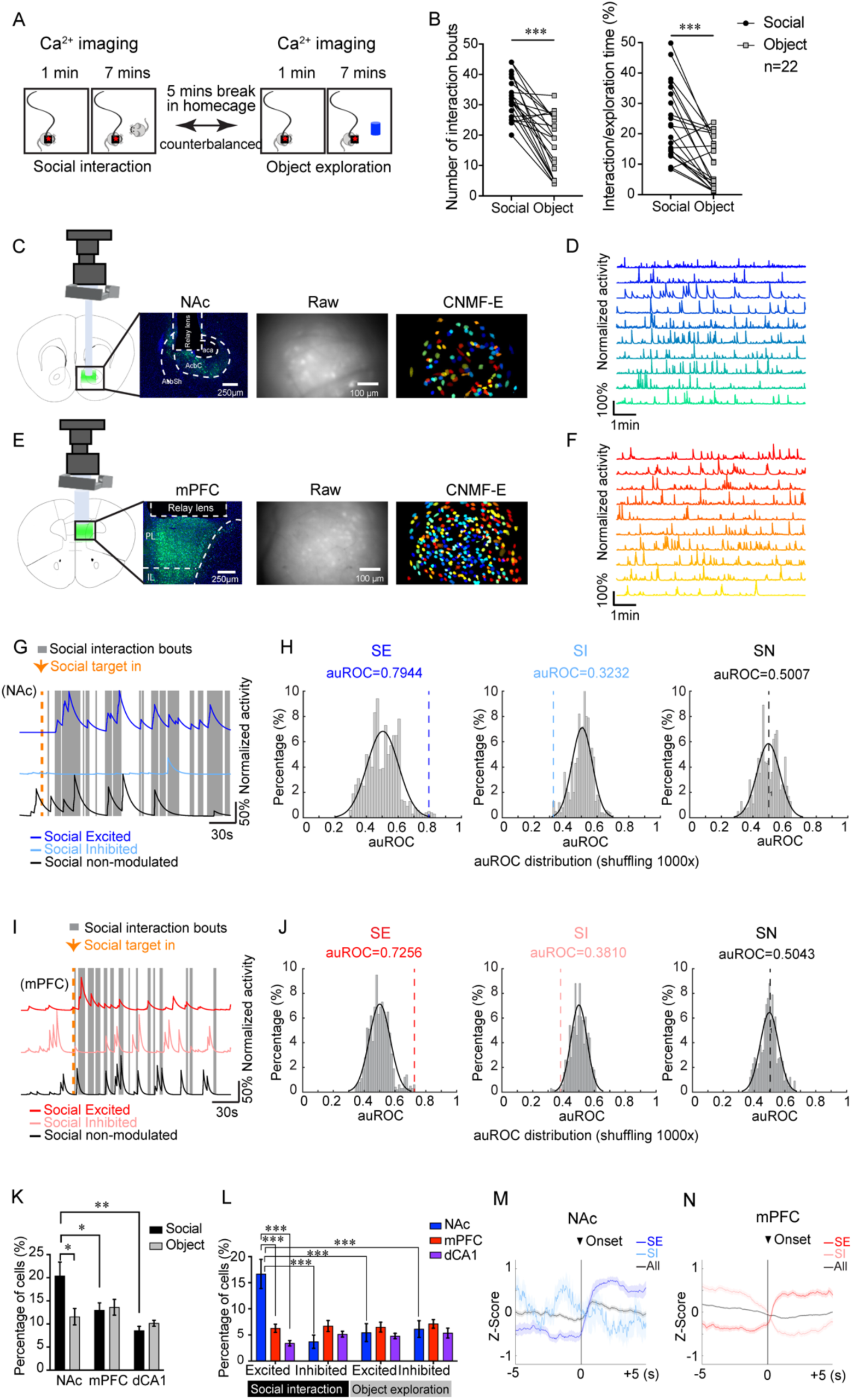
NAc, mPFC and dCA1 neural dynamics during social interaction and object exploration. (**A**) Schematic diagram of the social interaction assay counterbalanced with object exploration during calcium imaging with the miniature microscope. (**B**) Number of interaction bouts and the percentage of time the animal engaged in social interaction or object exploration. n=22 animals, including 11 recorded in NAc and 11 recorded in mPFC. Number of interaction bouts, Wilcoxon signed rank test, P<0.0001. Percentage of interaction/exploration time, Wilcoxon signed rank test, P<0.0001. (**C, E**), Left two columns: Schematic showing virus injection (Green, GCaMP6f; Blue, DAPI), GRIN lens implantation and miniscope placement for calcium imaging above NAc (core) and mPFC (PrL). Right two columns: raw calcium fluorescence of the field of view from example animals and single neurons extracted from the field of view by CNMF-E. (**D, F**) Example of calcium traces recorded in single sessions from NAc and mPFC. (**G, I**) Example calcium traces of social interaction-excited (SE), inhibited (SI), and non-modulated (SN) neurons in the NAc and mPFC characterized by ROC analysis and corresponding social interaction epochs. (**H, J**) Distribution of auROCs obtained from circular shuffling of calcium activity 1000 times. The auROC for the representative cells in G and I are indicated by the dash line. (**K**) Percentage of social interaction responsive neurons (including SE and SI) and object interaction responsive neurons (including OE and OI) in NAc, mPFC and dCA1. NAc, n=11 animals; mPFC, n=11 animals; dCA1 n=7 animals. For the comparison of percentage of social responsive cells in different brain regions and the comparison of percentage of object responsive cells in different brain regions, one-way ANOVA followed by *Holm*-*Šídák test*. Social cells, NAc vs mPFC, P=0.0421; NAc vs dCA1, P=0.0057. For the comparison of percentages of social responsive cells and object responsive cells within single brain regions, Wilcoxon signed rank test. NAc social cells vs object cells, P=0.0273. (**L**) Breakdown of social interaction/object exploration responsive cells (in K) into social excited/inhibited cells and object excited/inhibited cells. Two-way ANOVA followed by Tukey’s multiple comparisons test. (**M, N**) Average of responses of all cells, social excited cells, and social inhibited cells in the NAc and mPFC aligned to the onset of social interactions, presented in Z-score. Data shown as Mean ± SEM. *P<0.05, **P<0.01, ***P<0.001.

### NAc contains higher percentage of social cells than mPFC

The mPFC is also a critical hub for social behavior (*16–18*) and has projections to the NAc (*19*). To compare how the representation of social interactions differs between mPFC and NAc, we performed calcium imaging using miniaturized microscopy in the mPFC during social and object exploration (Fig. 1E) and did the same analysis that we had performed for NAc data (Fig. 1F, I, J). Although we could monitor a larger number of mPFC neurons (272.3±32.5) than NAc neurons (33.1±6.9) (Fig. S5A), the percentage of cells modulated by social interaction was significantly lower in mPFC than in NAc (Fig. 1K). Within mPFC, the percentages of social and object modulated neurons were similar (Fig. 1K, L) and the overlap between cells modulated by social interaction and those modulated by object exploration was 2.6±0.6% (Fig. S1A). As expected, in both NAc and mPFC, interaction onset-triggered averages of socially excited and inhibited neurons showed increases and decreases in fluorescence in response to the social interactions, respectively (Fig. 1M, N, object, Fig. S1B, C). Importantly, there was no significant correlation between the percentage of social interaction time and the percentage of social or object target responsive cells in both NAc and mPFC (Fig. S1D-G).

As a control, we recorded neural activity in dorsal CA1 (dCA1) (Fig. S2), a region well-known to be important in learning and memory (*20*, *21*), spatial information encoding (*22–24*), and object recognition processing (*25*, *26*), but not social interactions. In dCA1, a much smaller proportion of neurons was modulated by social interaction than in NAc (Fig.1K), and there was no difference between the percentages of social cells and object cells in dCA1 (Fig.1K, L).

mPFC neurons consist of a heterogeneous population of neurons with distinct projection targets which have been posited to drive distinct behaviors(*19*, *27–30*) through top-down control of information processing(*31*, *32*). To determine whether any of these mPFC projections play a privileged role in encoding social interactions, we used retrograde transported AAV expressing Cre and Cre-dependent GCaMP6f to label projection-specific mPFC neurons (Fig. S3A) and recorded their activities (Fig. S3B, C). We imaged mPFC-NAc, mPFC-mediodorsal thalamus (MDT), mPFC-ventral tegmental area (VTA), and mPFC-basolateral amygdala (BLA) neurons. Among these cell types, there was no statistically significant difference in the proportions of neurons modulated by social interaction (Fig. S3D), though proportions of neurons engaged in social interaction in these projections were lower than in the NAc.

To investigate the possibility that the neural populations we recorded also encode velocity, we used DeepLabCut (*33*) to track the subject mouse and calculate its velocity during social and object interactions. Compared to shuffled control data (Fig. S4A), although percentages of neurons significantly modulated by velocity in NAc and mPFC were similar (12.7±5.0%, 12.3±0.8%) (Fig. S4B, C), only in mPFC was this proportion significantly greater than chance (Fig. S4B). Importantly, only 21.1±7.1% of socially modulated neurons in NAc also encoded velocity and this overlap was not significantly different from chance (Fig. S4F). Therefore, social representations do not simply arise from the locomotor activity of animals (Fig. S4D-G).

### Specificity of NAc neural population encoding social interaction

How do population activity patterns in NAc encode social interaction? To answer this question, we built a decoder based on partial least squares regression (*34*). We used ROC analysis to evaluate the decoder performance and compared the performance based on recorded data with the performance based on shuffled data. Interestingly, we found that decoders trained on NAc population neural activity patterns could predict social interaction epochs but not object exploration epochs (Fig. 2A, B). On the other hand, decoders trained on mPFC population neural activity could predict epochs of both social interaction and object exploration (Fig. 2C, D). As expected, the decoder trained with dCA1 neural population activity could not predict social interaction epochs but could predict object exploration epochs with high reliability (Fig. 2E, F).

**Fig. 2.**
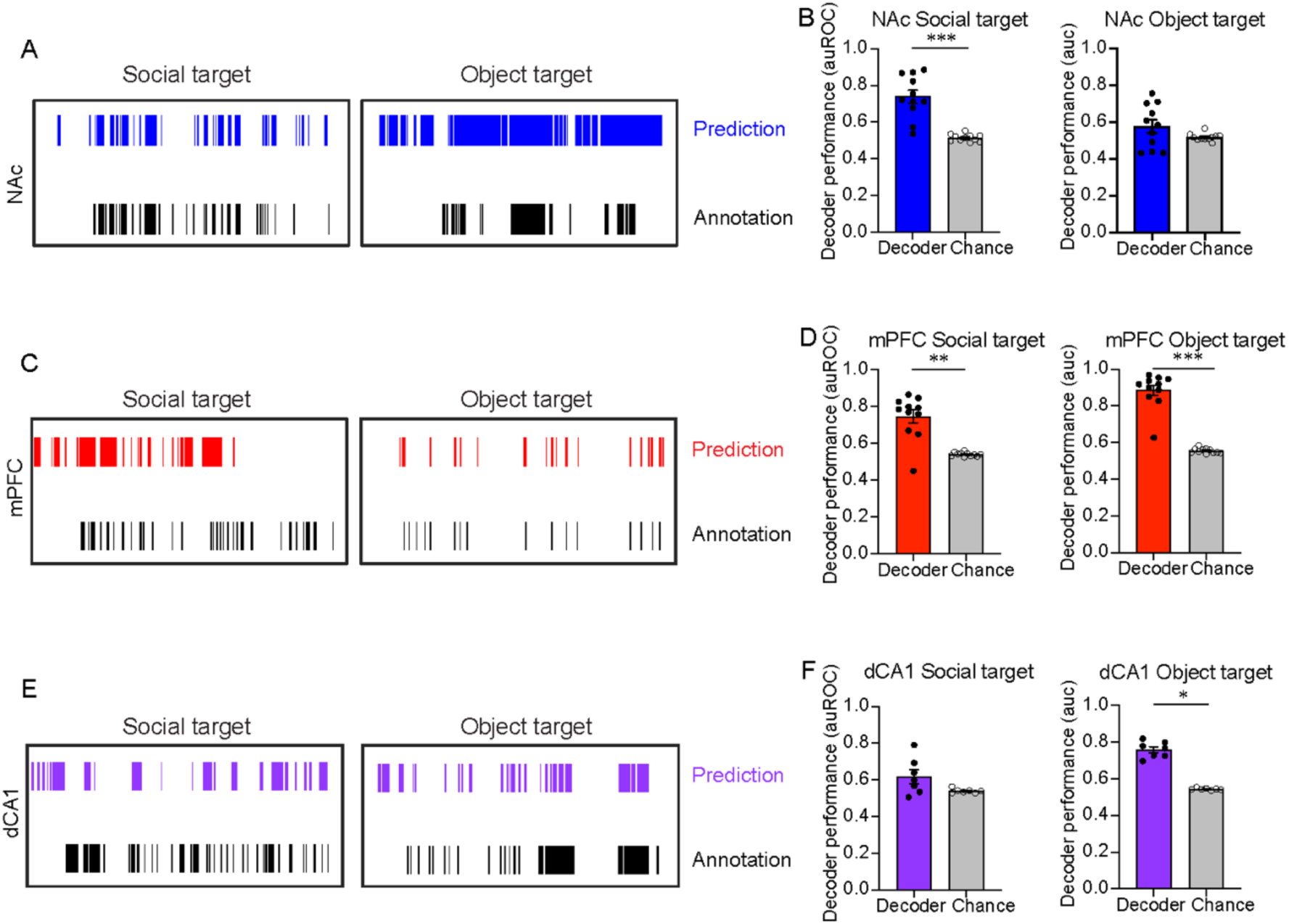
NAc neural population specifically encodes social interaction. (**A, C, E**) Examples of predicted epochs of social interaction and object exploration with partial least squares regression performed on population activity recorded from NAc, mPFC and dCA1. (**B, D, F**) Decoder performance (auROC) predicting social interaction and object exploration with NAc, mPFC and dCA1 population neural activities. Only NAc population activity can specifically predict interaction epochs with the social target. In B, n=11 animals, Wilcoxon signed rank test, social, P=0.0010, object, P=0.1016. In D, n=11 animals, Wilcoxon signed rank test, social, P=0.002, object, P=0.0010. In F, n=7 animals, Wilcoxon signed rank test, social, P=0.1562, object P=0.0156. *P<0.05, **P<0.01, ***P<0.001.

Due to technical limitations, the total numbers of cells we imaged from different regions differed significantly (Fig. S5A). To exclude the possibility that cell numbers might affect decoder performance, we randomly chose a similar number of cells (30 cells) from mPFC and dCA1 and repeated decoding analysis. The results of this analysis with equal cell numbers were similar to our previous results (Fig. 2, Fig. S5B, C). Moreover, when we compared decoder performance trained with a comparable number of cells among the three brain regions, the NAc population neural activity was significantly better at predicting social interaction epochs than mPFC and dCA1 population neural activity (Fig. S5D, left panel). Therefore, even small populations of NAc neurons can reliably encode social interaction.

### NAc D1-MSNs preferentially encode social interactions

NAc receives and integrates cognitive and affective information from upstream brain regions in a cell type specific manner (*4*, *5*, *10*, *11*, *19*, *35*, *36*). Previous studies have indicated that two major MSNs in NAc, D1-MSNs and D2-MSNs may drive distinct behaviors (*6–9*). Whether these distinct cell types differentially represent social interactions in NAc is still not understood (*10*, *11*). To solve this question, we compared the neural dynamics of D1- and D2-MSNs in the NAc core region during social and object interactions in D1- and A2A (a D2-MSN-specific Cre line)- Cre mice lines (Fig. 3A). There were more neurons tuned to social interaction than neurons tuned to object exploration in D1-Cre mice, while there was no difference in the proportion of these neurons in A2A-Cre mice (Fig. 3B). Specifically, the proportion of social excited cells was significantly larger in D1-Cre mice than in A2A-Cre mice (Fig. 3C). Furthermore, while a PLSR decoder trained with D1-MSN neural activity could specifically predict social interaction, but not object exploration (Fig. 3D, E), the decoder trained with D2-MSN neural activity could predict neither social interaction nor object exploration (Fig. 3F, G). Therefore, D1-MSNs in NAc core are the main cell type representing social interaction.

**Fig. 3.**
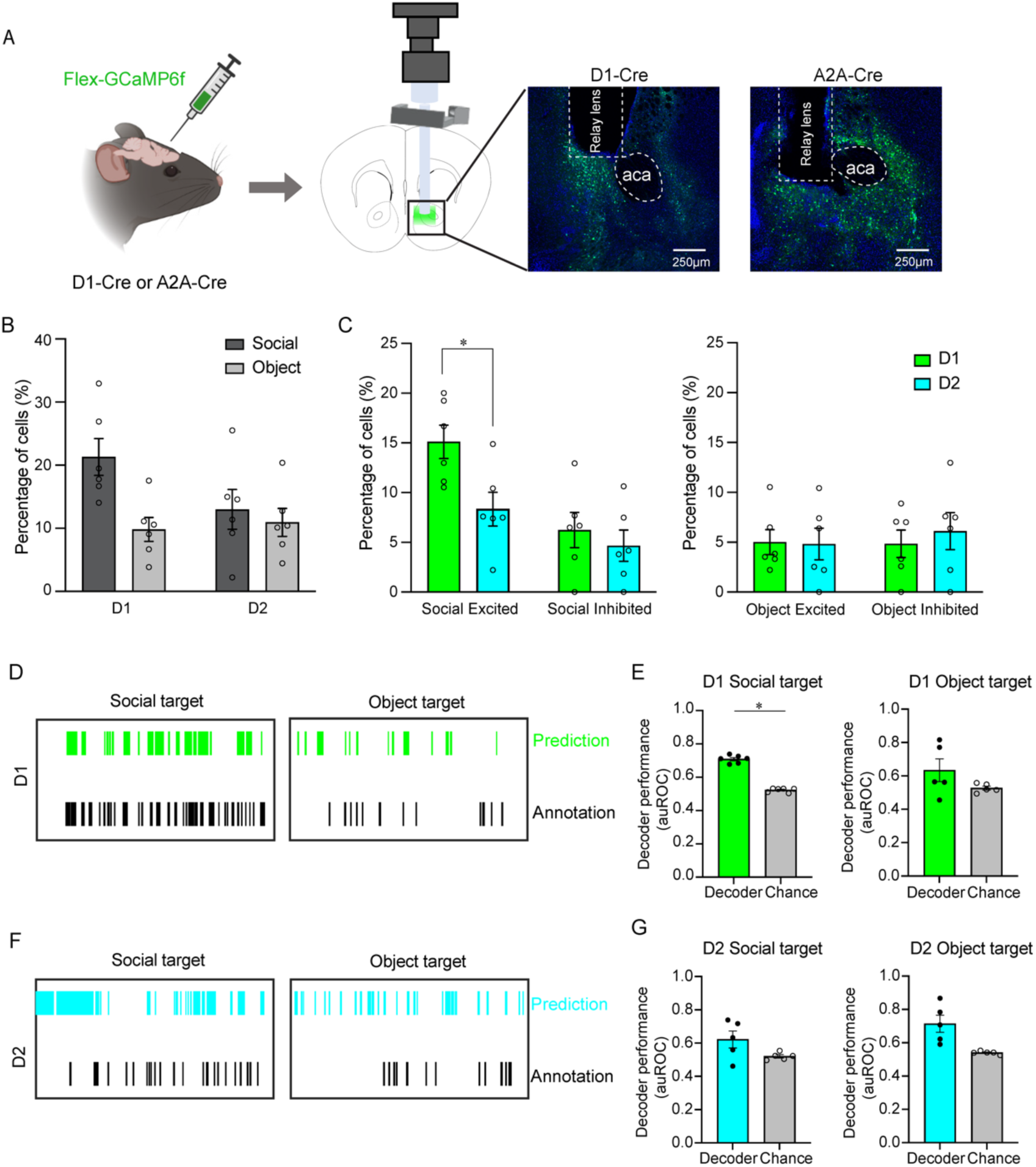
Neural representation of social interaction by D1R-expressing MSNs (D1-MSNs) in NAc. (**A**) Diagram of labelling D1-MSNs and D2-MSNs and lens implanting in D1-Cre and A2A-Cre mice. (**B**) Percentage of social interaction (SE+SI) and object exploration (OE+OI) responsive cells in D1-Cre and A2A-Cre mice. n=6 animals for D1-Cre mice and n=6 animals for A2A-Cre mice. Wilcoxon signed rank test. D1-Cre mice, P=0.065; A2A-Cre mice, P=0.5625. (**C**) Percentage of D1-MSNs and D2-MSNs in NAc excited/inhibited by social interaction and object exploration. n=6 animals for D1-Cre mice and n=6 animals for A2A-Cre mice. Mann-Whitney test. Social excited, P=0.0152; social inhibited, P=0.6234; object excited, P=0.7338; object inhibited, P=0.7381. (**D, F**) Examples of predicted epochs of social interaction and object exploration using population activity recorded from D1- or D2-MSNs in the NAc of D1-Cre and A2A-Cre mice. (**E, G**) Performance of decoders trained on D1-MSNs’ and D2-MSNs’ neural activities. Population activity of D1-MSNs in the NAc can predict social interaction epochs. Wilcoxon signed rank test. E, n=6 animals for social interaction decoding. P=0.0312. n=5 animals for object exploration decoding. P=0.3125. G, n=5 animals for both social interaction and object exploration decoding; Social, P=0.1875; Object, P=0.0625. *P<0.05.

### Stable NAc neural representation of social interaction

How is social information encoded longitudinally in NAc and mPFC? To answer this question, we performed calcium imaging of the same neuronal populations while mice socially interacted across 6 days (Fig. 4A). In this task, the imaged mouse interacted with the same social target during each session. Although the imaged mouse spent a decreasing percentage of time interacting with the other mouse across the total time within each individual session (Fig. S6A), the percentage of time spent interacting with the other animal did not change significantly over days (Fig. S6B). Using CellReg (*37*), we tracked the same neurons across six days (Fig. 4A-C). After registering neurons across all sessions of calcium imaging, we calculated the proportion of active neurons on Day 1, that were active anytime during the session (not only during epochs of social interaction) on later days (Day 2 to Day 6). 45.6± 2.5% and 70.7± 1.1% of neurons that were active on Day 1, were active on the subsequent days of imaging in NAc and mPFC, respectively (Fig. 4D). Similarly, the proportion of neurons that were active on all six days of imaging was significantly lower in NAc than in mPFC (Fig. S6C). The proportion of socially modulated neurons (SE+SI) that continued to be socially modulated on subsequent days (overlap of socially modulated neurons between Day 1 and Day X) was nearly 30% in NAc (Day1 ∩ Day2) and declined to 15-20% on subsequent days. This proportion of socially modulated neurons was not significantly different in mPFC (Fig. 4E).

**Fig. 4.**
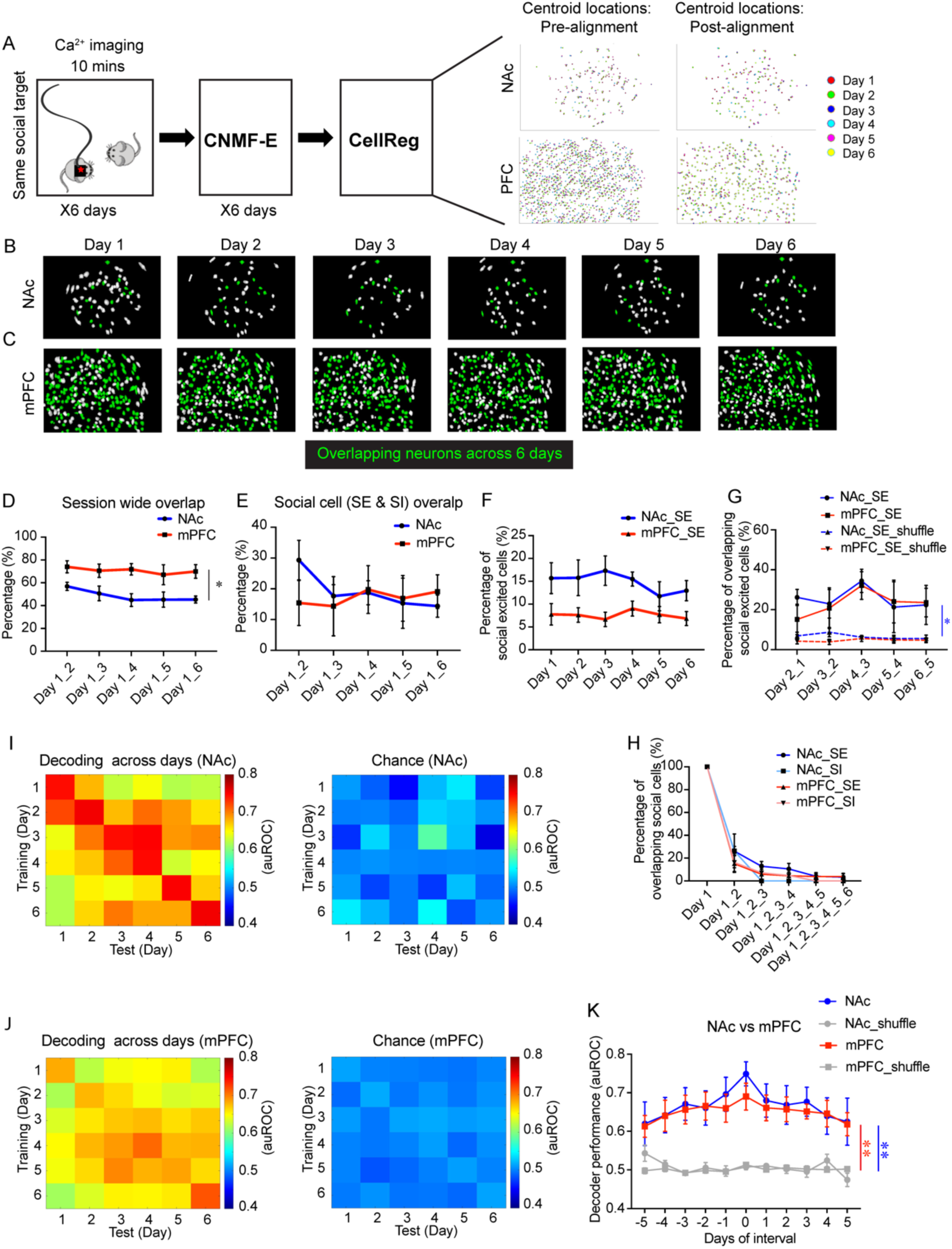
Stability analysis of long-term neural representations of social interaction. (**A**) Schematic diagram of social interaction assay with miniscope calcium imaging across six days (same social target across multiple sessions). Calcium traces were extracted by CNMF-E. The same neurons across days were tracked by CellReg. (**B**, **C**) Examples of cell registration across six days using the CellReg package. Green, active cells during social interaction every day. (**D**) Session-wide overlap of active cells on each day (refer to Day 1) in NAc and mPFC. NAc, n=5 animals; mPFC, n=4 animals. Two-way repeated-measures ANOVA, P=0.0199. (**E**) Overlap of social cells (SE & SI) on each day (refer to Day 1) in NAc and mPFC. Two-way repeated-measures ANOVA, P=0.8013. (**F**) Percentage of social excited cells in NAc and mPFC on each day. Two-way repeated-measures ANOVA, P=0.0502. (**G**) Overlap of social excited cells in NAc and mPFC between two consecutive days. Two-way repeated-measures ANOVA, NAc_SE vs NAc_SE_shuffle, P=0.0218; mPFC_SE vs mPFC_SE_shuffle, P=0.0550. (**H**) Overlap of social excited/inhibited cells in NAc and mPFC across different number of days. (**I, J**) Matrix showing decoder performance of cross days prediction of social interaction using NAc and mPFC population neural activities. (**K**) Plot of decoder performance in I and J. Two-way repeated-measures ANOVA, NAc vs NAc_shuffle, P=0.0059; mPFC vs mPFC_shuffle, P=0.0032. *P<0.05, **P<0.01.

We then analyzed the stability of socially excited and inhibited neurons separately. Consistent with data obtained from one-session social interaction tests, during long-term social interaction tests, the proportion of neurons socially excited was generally higher on each day in NAc compared to mPFC (though this did not reach significance due to multiple-comparisons) (Fig. 4F). There was no difference between percentages of social inhibited cells in NAc and mPFC (Fig. S6D). We examined the proportion of socially excited neurons that were still socially excited on the next day of imaging (overlap between Day X and Day X+1). This proportion was around 25% for both NAc and mPFC, and higher than chance level of reactivation in NAc (Fig. 4G). The proportion of social inhibited cells was similar in NAc and mPFC, and higher than chance level in NAc (Fig. S6E). We subsequently examined the proportion of neurons that were similarly excited or similarly inhibited on any pairs of days during the six-day recording period. Interestingly, there was a similar overlap of socially excited or inhibited neurons regardless of the gap between imaging days (Fig. S6G, H). The probability that a socially excited or socially inhibited neuron imaged on Day 1 would remain socially excited or socially inhibited for 2, 3, 4, 5 or 6 days progressively diminished such that fewer than 4% of all neurons were consistently socially excited and no cells were consistently socially inhibited during all six days (Fig. 4H, Fig. S6F, right panel), mostly due to a lower initial total percentage of social inhibited cells, especially for NAc. Taken together, these results suggest a high turnover of neurons responsive to social interaction. While an equal proportion of neurons represented social interaction on each day (Fig. 4F), single neurons joined or left the ensemble on any day.

It is important to note that the socially modulated neurons (SE+SI) analyzed here were only those showing significant response to social interaction (Fig. 1G-J). What were responses of neurons like during sessions when they were identified as *not* being significantly modulated by social interaction? Though not passing the significance threshold, was there a tendency for neurons to respond to social interaction in a consistent manner, either positively (excitatory response) or negatively (inhibitory response)? We first plotted an auROC (ROC analysis result for each neuron) map of each mouse listing all cells detected across six days of recordings (Fig. S6I, J). We then calculated the percentage of cells that tended to positively or negatively respond to social interaction by counting cells which had been detected at least during two recording sessions and showed a consistent tendency (with auROC always >0.5 or auROC always <0.5). We found that the NAc region contained a higher percentage of cells consistently positively modulated by social interaction when compared with mPFC (Fig. S6K).

Despite the large-scale changes in the response properties of individual neurons, mouse behavior (percentage of interaction time on each day) did not change. Therefore, population activity patterns may still allow stable encoding of social interactions. We performed cross-day decoding of social interaction epochs by training the PLSR decoder with neural activity patterns from any of the 6 recordings days (Day X, row in matrix), applied it to predict social interactions on all other days (Day Y, column in matrix), and repeated this process with shuffled activity data (Fig. 4I, J). By comparing the decoder performance matrix derived from real data with that derived from shuffled data and linearizing the matrices into vectors (Fig. S6L), the decodability for each combination of days was found to be greater than shuffled data. Additionally, we calculated the performance of decoders with different lengths of time separating training and testing epochs, which was significantly better than performance of the decoders with shuffled data (Fig. 4K). mPFC population decoding showed similar results (Fig. 4J, K, Fig. S6M), though same-day decoding, as shown by the diagonal in the decoding matrix, was generally worse in mPFC than in NAc (Fig. 4I, J). Taken together, our results suggest that although the majority of neurons modulated by social interaction on one day are no longer socially modulated in subsequent days, the populational representation of social interactions remains stable across the recording period.

### Distinct ensembles encode rewards derived from social interaction and sucrose consumption

We have shown that a sizable proportion of NAc neurons encode social interaction, and that this representation is stable across days. Nonetheless, it is not yet known whether social interactions are encoded distinctly from other types of rewards such as sucrose consumption. Do distinct populations carry information about the different rewards? Rewarding stimuli (other than social reward) have been shown to be represented in the NAc (*38–40*). These include natural rewards derived from foods such as sucrose. Previous studies have shown ensembles activated by sucrose consumption to be distinct from ensembles activated by cocaine intake in the NAc core, suggesting that distinct types of rewards may be represented by different ensembles (*38*, *40*). We therefore trained mice to lick a spout to obtain a 10% sucrose solution. We then imaged NAc or mPFC during 10-minute sessions of sucrose consumption or social interaction, separated by 60 minutes (Fig. 5A). We performed ROC analysis to identify neurons responding to social interaction epochs or sucrose consumption. In NAc, 24.6 ± 3.2% of neurons were modulated by social interactions and 30.9 ± 4.0% by sucrose consumption (Fig. 5B). In mPFC, 16.9 ± 2.3% of neurons were modulated by social interactions and about 27.5 ± 2.2% by sucrose consumption (Fig. 5C). The overlap between sucrose and socially modulated neurons was similar to that expected by chance in both NAc and mPFC (Fig. 5D, E). Social and sucrose modulated neurons were intermingled in both NAc and mPFC and did not show any evidence of clustering (Fig. S7). Therefore, distinct intermixed ensembles in NAc and mPFC encode rewards derived from social interactions and sucrose consumption.

**Fig. 5.**
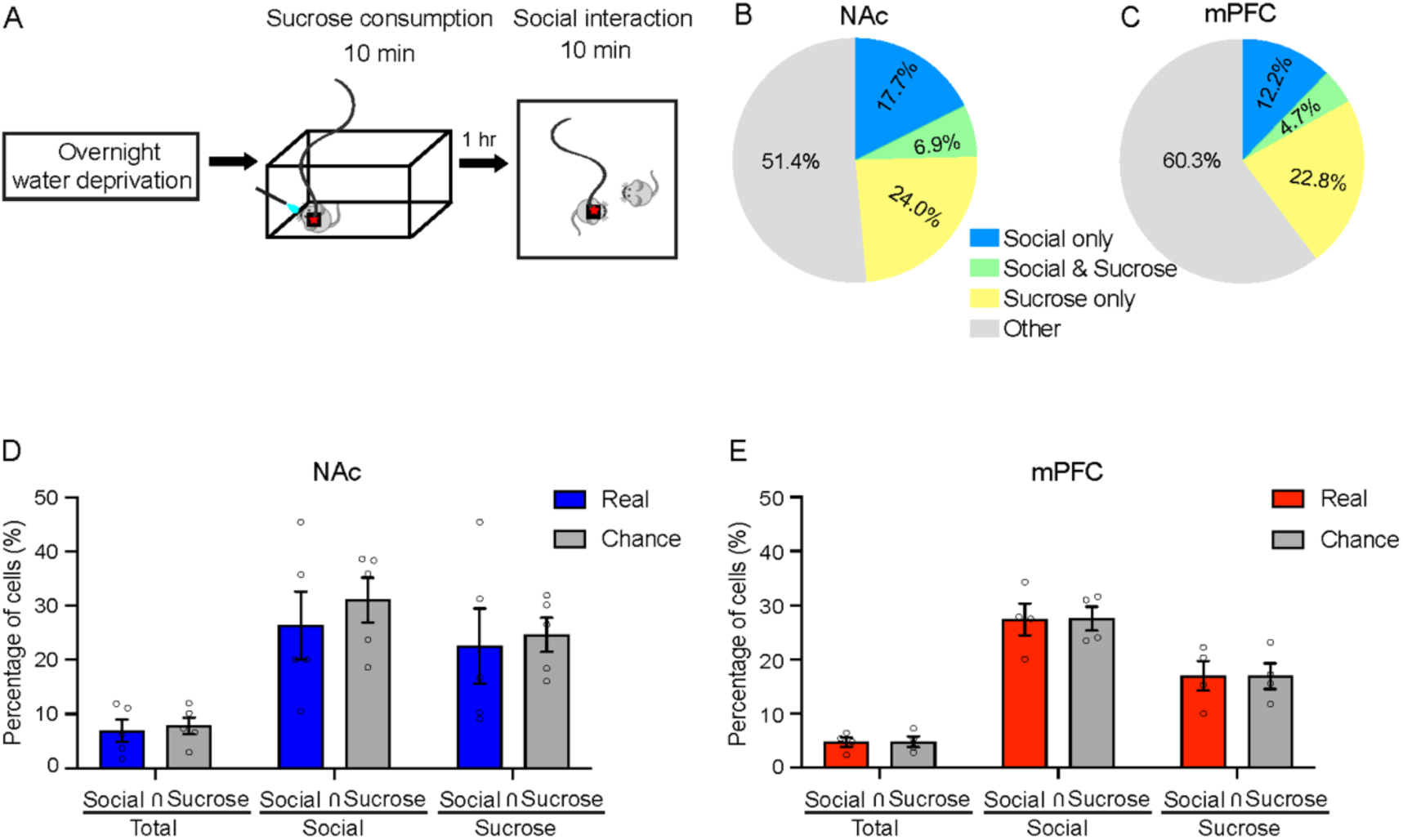
Distinct ensembles encode social interaction and sucrose consumption. (**A**) Schematic of behavior assay of sucrose consumption and social interaction. (**B, C**) Percentages of cells in NAc and mPFC responding to sucrose consumption and/or social interaction. NAc, n=5 animals; mPFC, n=4 animals. (**D, E**) Percentages of cells encoding both social interaction and sucrose consumption over total cells, social interaction tuned cells and sucrose consumption tuned cells compared with chance level. Two-way ANOVA followed by Fisher’s LSD test, P>0.05 for all.

### Social representation deficits in Cntnap2 knockout mice

If the stable social representations of NAc are important for driving social interactions, then one would expect these representations may be disrupted in a mouse model of ASD with diminished social interactions. We therefore performed calcium imaging during social interactions in NAc of Cntnap2^-/-^ (KO) mice and their littermate controls. The Cntnap2^-/-^ model has high face and construct validity; nearly two thirds of the children with Cntnap2 homozygous loss-of-function mutations are diagnosed with ASD (*41*). In addition, Cntnap2 KO mice have deficits in social behavior and exhibit repetitive behaviors, hyperactivity, and epileptic seizures (*42–44*). We compared the proportion of time subject male mice spent interacting with a male C57/BL6J mouse versus interacting with an object during separate 8-minute sessions (same as Fig. 1A). Both KO and wildtype (WT) mice spent a greater proportion of time interacting with the other mouse than exploring the object (Fig. 6A). However, KO mice spent less time interacting with the other mice from the second minute of the social interaction session onward compared to WT mice (Fig. 6B). The mean duration of each social interaction epoch was shorter in KO mice compared to WT mice (Fig. 6C). These results suggest that KO mice lost interest in the social target earlier, possibly because social interactions were less rewarding for KO mice. Compared to KO mice, there was a trend of a smaller proportion of social interaction modulated neurons in KO mice (SE+SI, 31.46± 6.68% from 6 WT mice, 18.58±3.28%, mean ± SEM, from 9 KO mice), though this difference did not reach statistical significance (Fig. 6D). The proportions of object modulated neurons identified in NAc in WT and KO mice are similar (OE+OI, 17.02± 3.68% from 6 WT mice, 18.61± 2.41%, mean ± SEM, from 9 KO mice) (Fig. S8A). When we quantified the magnitude of social interaction evoked activity in WT and KO mice by calculating the area under curve (activity AUC) we found that NAc social excited neurons in KO mice showed significantly lower AUC values, suggesting smaller socially evoked responses (Fig. 6E, F Fig. S8B-D). As the Cntnap2^-/-^ mice lost interest in social interactions faster than WT mice, we hypothesized that social responses in KO NAc neurons would adapt more vigorously than in WT mice. We plotted the magnitude of the first five NAc calcium responses elicited for each social interaction epoch in KO and WT mice. The magnitude of responses to the first five social interaction epochs decayed to baseline sooner in Cntnap2^-/-^ animals, and the response to each epoch is weaker than in WT animals starting from the first epoch, suggesting more rapid adaptation of NAc responses in KO mice, though there was a great variability in the dynamics of responses (Fig. S8E).

**Fig. 6.**
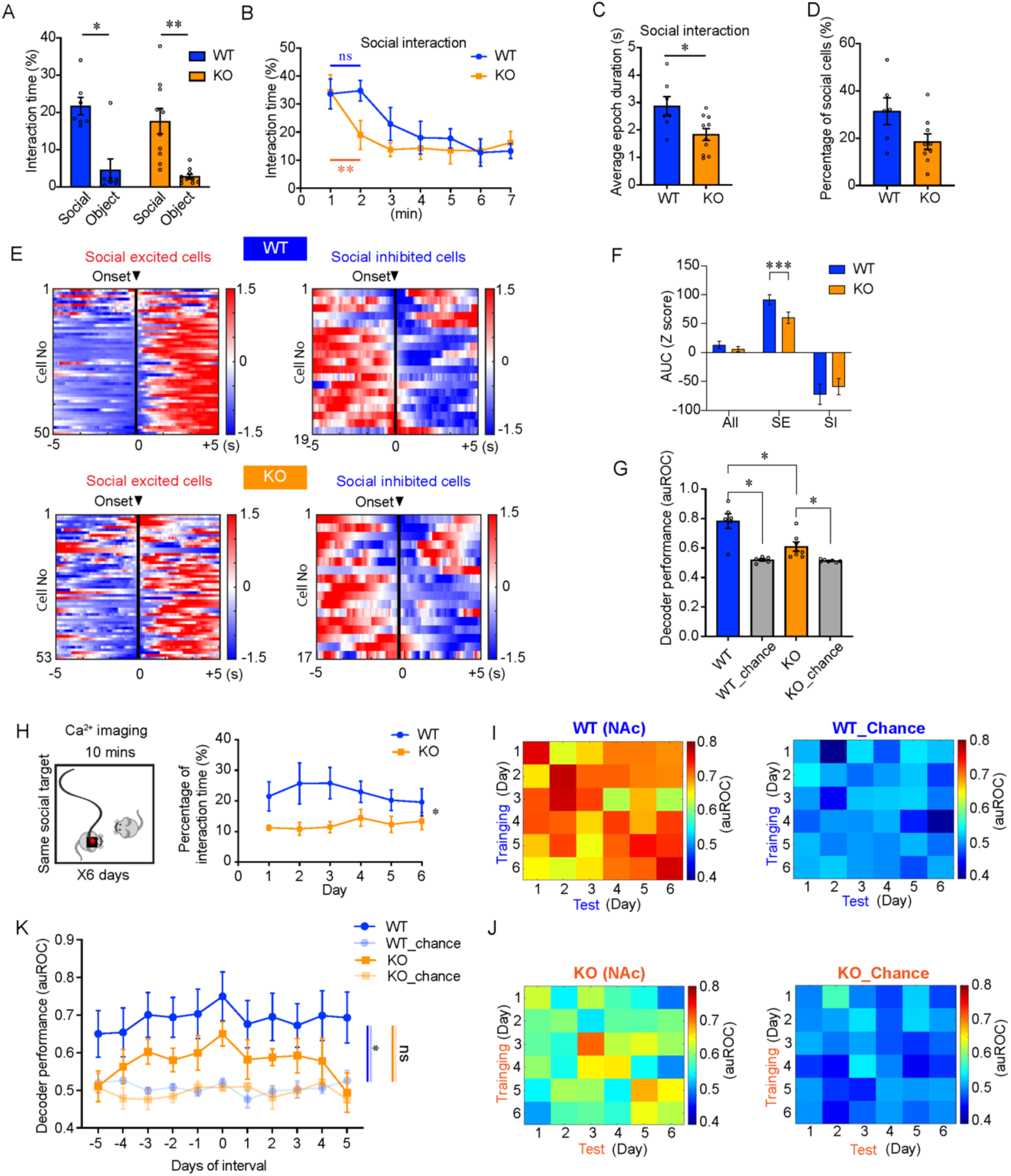
Degraded NAc social representation of Cntnap2^-/-^ mice. (**A**) Percentage of time spent on social interaction and object exploration by control and Cntnap2^-/-^ mice. WT, n=7 animals; KO, n=10 animals. Wilcoxon signed rank test, WT, P=0.032; KO, P=0.002. (**B**) Percentage of social interaction time during each minute of behavior test. WT, n=7 animals; KO, n=10 animals. One-way repeated-measures ANOVA. WT 1^st^ second vs 2^nd^ second, P=0.8349; KO 1^st^ second vs 2^nd^ second, P=0.0062. (**C**) Average duration of each interaction epoch. WT, n=7 animals; KO, n=10 animals. Mann-Whitney test. P=0.033. (**D**) Percentage of social interaction modulated cells identified in WT and KO mice in NAc. WT, n=6, animals; KO, n=9 animals. Mann-Whitney test. P=0.0931. (**E**) Average responses of social excited and inhibited cells to the onset of social interaction epochs. WT, n=6 animals; KO, n=9 animals. (**F**) Statistical analysis of activity change in WT and KO aligned by onset of social interaction, shown by area under curve value (AUC, area post the onset of social interaction) and Z-score. Mann-Whitney test. All neurons: WT, n=211 neurons; KO, n=375 neurons. P=0.1901. SE: WT, n=50 neurons; KO, n=53 neurons. P=0.0008. SI: WT, n=19 neurons; KO, n=17 neurons. P=0.2711. (G) Decoder performance predicting single session social interaction with WT and KO NAc neural activities compared to chance level. WT, n=6 animals; KO, n=7 animals. Wilcoxon signed rank test, WT vs WT_chance, P=0.0312; KO vs KO_chance, P=0.0156. Mann-Whitney test, WT vs KO, P=0.0221. (**H**) Percentage of social interaction time spent by WT and KO mice during long-term social interaction assay. WT, n=6 animals; KO, n=7 animals. Two-way repeated-measures ANOVA. P=0.0388. (**I, J**) Long-term social representation of WT and KO NAc indicated by the performance of cross-day decoding of social interaction compared to chance level. (**K**) Plot of decoder performance in I and J. WT, n=4 animals; KO, n=4 animals. Two-way repeated-measures ANOVA, WT vs WT_shuffle, P=0.0187; KO vs KO_shuffle, P=0.0721. *P<0.05, **P<0.01, ***P<0.001, ns, not significant.

Are population representations of social interactions degraded in KO mice? We trained a PLSR decoder with either KO or littermate control NAc calcium imaging data during the social interaction test and found that, while both WT and KO NAc activity could predict social interactions at accuracy levels greater than chance, decoders performed at higher accuracy when trained and tested on data from WT mice compared to KO animals (Fig. 6G).

To examine long-term social representation in Cntnap2^-/-^ mice, we performed calcium imaging in NAc during six days of social interaction with the same intruder (as described in Fig. 4) (Fig. 6H, left panel) during 10-minute sessions. Generally, WT mice spent more time interacting with the other mouse in the same arena on each day (Fig. 6H, right panel). We then analyzed the social representation consistency of the neurons recorded from NAc as described in Fig. S6I-K. NAc neurons in KO mice showed a lower consistency in positively responding to social interaction when compared to WT mice (Fig. S9A). We then plotted those neurons that were present across six days and sorted them by their auROC value on the first day (Fig. S9B, C), which showed lower representational consistency of NAc neurons and lower response strength in KO mice.

We then conducted cross-day decoding using each day’s neural activity pattern to train the decoder and then predicted social interaction epochs during that day and every other day (Fig. 6I, J). Consistent with the one session test decoding result (Fig. 6G), KO mice showed worse same-day decoding results (diagonal line in matrix) compared to WT mice. WT mice showed significantly higher cross-day decoding performance compared to chance level. However, cross-day decoding of social interactions in Cntnap2^-/-^ mice was significantly degraded (Fig. 6K, Fig. S9D, E). Decoders trained from NAc dynamics in KO mice could not predict social interactions across days as well as those trained from NAc dynamics in WT mice, indicating that social behavioral deficits were accompanied by long-term social representation deficits in NAc.

### Cell type specific control of social interactions by NAc neurons

To further dissect the neural mechanism driving social interaction representations, we firstly non- specifically inhibited all neurons in NAc core by stimulating NpHR3.0 driven by human synapsin promoter during social interaction and object exploration (Fig. 7A). We compared the percentage of social interaction time between the NpHR group and EGFP group during light ON sessions; we also compared the percentage of social interaction time in the light ON and light OFF sessions in the NpHR group. Surprisingly, non-specific NAc core inhibition significantly increased social interaction time (Fig. 7B, C) with no effect on object exploration (Fig. S10A). Since mice were most likely to interact with other mice during the first two minutes of the interaction session (Fig. 6B), we analyzed the percentage of interaction time during first two-minute block (minute 1 and 2) and second two-minute block (minute 3 and 4). Non-specific NAc core inhibition resulted increased interaction time during both time periods (Fig. 7D, E). There was a larger increase in the interaction time during the second two-minute block than that during first 2-minute block (Fig. 7E). In addition, NpHR inhibition of neurons prolonged the average interaction epoch (Fig. S10E).

**Fig. 7.**
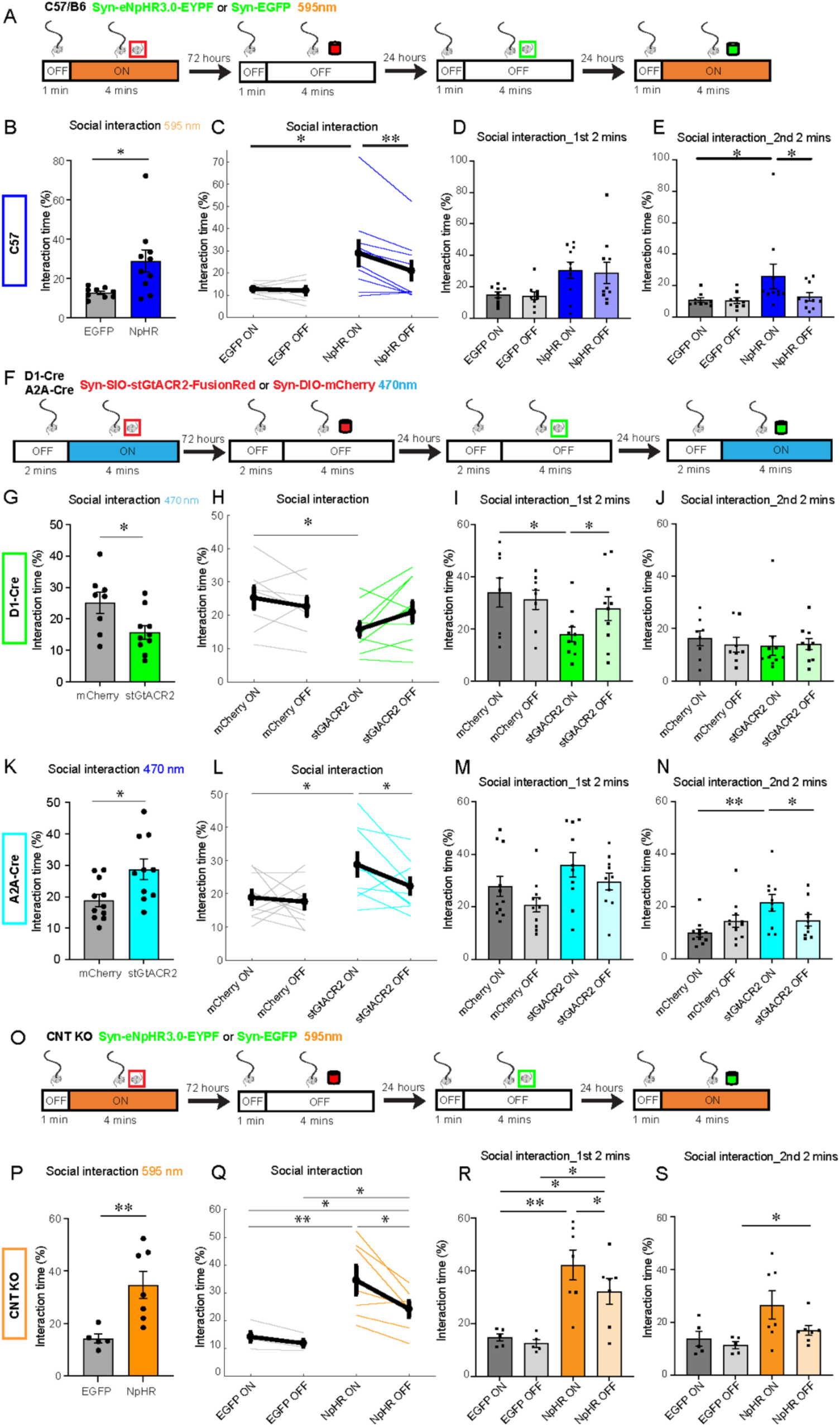
Selective optogenetic manipulation and rescue of social deficit in Cntnap2^-/-^ mice. (**A, F, O**) Behavioral paradigm of NpHR inhibition of NAc core in C57 mice (A), selective stGtACR2 inhibition of NAc D1-MSNs and D2-MSNs in D1-Cre and A2A-Cre mice (F), and NpHR inhibition of NAc core neurons in Cntnap2^-/-^ mice (O). (**B, C, G, H, K, L, P, Q**) Percentage of social interaction time during each session. Mann-Whitney test for comparing interaction time of mice with different opsins. Wilcoxon signed rank test for comparing the same mice under different condition. (B) C57 mice, EGFP, n=9 animals; NpHR, n=10 animals. EGFP light ON vs NpHR light ON, P=0.0101. (C) NpHR light ON vs NpHR light OFF, P=0.0098. (G) D1-Cre mice, mCherry, n=8 animals; stGtACR2, n=10 animals. mCherry light ON vs stGtACR2 light ON, P=0.0434. (H) stGtACR2 light ON vs stGtACR2 light OFF, P=0.1055. (K) A2A-Cre mice, mCherry, n=11 animals; stGtACR2, n=10 animals. mCherry light ON vs stGtACR2 light ON, P=0.0197. (L) stGtACR2 light ON vs stGtACR2 light OFF, P=0.0371. (P) Cntnap2^-/-^ mice, EGFP, n=5 animals; NpHR, n=7 animals. EGFP light ON vs NpHR light ON, P=0.0051. (Q) stGtACR2 light ON vs stGtACR2 light OFF, P=0.0312. EGFP light ON vs NpHR light OFF, P=0.048; EGFP light OFF vs NpHR light OFF, P=0.0101. (**D, E, I, J, M, N, R, S**) Percentage of social interaction time during the first and the second two-minute blocks. Mann-Whitney test for comparing interaction time of mice with different opsins. Wilcoxon signed rank test for comparing the same mice under different condition. (D) EGFP light ON vs NpHR light ON, P=0.0535; NpHR light ON vs NpHR light OFF, P=0.8457. (E) EGFP light ON vs NpHR light ON, P=0.0133; NpHR light ON vs NpHR light OFF, P=0.0273. (I) mCherry light ON vs stGtACR2 light ON, P=0.0434; stGtACR2 light ON vs stGtACR2 light OFF, P=0.0137. (J) mCherry light ON vs stGtACR2 light ON, P=0.2641; stGtACR2 light ON vs stGtACR2 light OFF, P=0.3223. (M) mCherry light ON vs stGtACR2 light ON, P=0.1517; stGtACR2 light ON vs stGtACR2 light OFF, P=0.2207. (N) mCherry light ON vs stGtACR2 light ON, P=0.0037; stGtACR2 light ON vs stGtACR2 light OFF, P=0.037. (R) EGFP light ON vs NpHR light ON, P=0.0025; NpHR light ON vs NpHR light OFF, P=0.0312; EGFP light ON vs NpHR light OFF, P=0.0177; EGFP light OFF vs NpHR light OFF, P=0.0101. (S) EGFP light ON vs NpHR light ON, P=0.1061; NpHR light ON vs NpHR light OFF, P=0.0781; EGFP light ON vs NpHR light OFF, P=0.4318; EGFP light OFF vs NpHR light OFF, P=0.0481. *P<0.05, **P<0.01.

Although our calcium imaging data indicated that D1-MSNs were more involved in encoding social interactions than D2-MSNs (Fig. 3), D2-MSNs may still regulate social interaction in a distinct manner. We therefore selectively inhibited either D1-MSNs or D2-MSNs in NAc core using Cre-dependent soma-targeted GtACR2 (stGtACR2) (Fig. 7F). Selectively inhibiting D1-MSNs decreased social interaction time (Fig. 7G, H) and average epoch duration (Fig. S10F). This effect was mostly observed during the first two minutes when mice showed greater interest in interacting with the social target (Fig. 6B, Fig. 7I, J). Conversely, inhibition of D2-MSNs increased the percentage of social interaction time (Fig. 7K, L) as well as average interaction epoch duration (Fig. S10G), mirroring what was seen with non-specific NAc inhibition. This significant increase in interactions was observed mostly during the second two minutes of the test session when mice typically decreased their social interactions (Fig. 6B, Fig. 7M, N). Inhibition of either D1- or D2-MSNs did not affect object exploration (Fig. S10B, C). Therefore, D1 and D2-MSNs played reciprocal roles in modulating social behavior. As non-specific inhibition of all NAc core neurons showed a similar effect to that observed with D2-MSN specific inhibition, D1 and D2-MSN activity may be differentially weighted in their ability to control social behavior, with D2 activity over-riding D1 activity.

Finally, we asked if non-specific inhibition of NAc neurons could rescue social deficit in Cntnap2^-/-^ mice (Fig. 7O). Non-specific NAc core neuron inhibition in Cntnap2^-/-^ mice significantly increased social interaction time (Fig. 7P, Q) and average epoch duration (Fig. S10H) without any effect on object exploration (Fig. S10D). Significant increase in social interaction time occurred during the first two minutes of the test session during which KO mice showed less social interaction than controls (Fig. 7R, S). Intriguingly, our experiments with NpHR or stGTACR2 showed that the effect of NAc inhibition or D2-neuron inhibition may have outlasted the optogenetic inhibition period by at least 96 hours, as mice expressing inhibitory opsins under light OFF condition (96 hours after the light ON session) showed a higher percentage of interaction time than mice which only expressed EGFP group or mCherry (Fig. 7C, L, Q). Therefore, inhibition of NAc core neurons can reverse the social deficits in the Cntnap2 model of autism.

## Discussion

Using miniscope calcium imaging (*20*, *45*, *46*), our study showed that NAc had a higher proportion of neurons activated by social interaction than mPFC or CA1 and could decode social interactions more efficiently. D1-MSNs in NAc were the main cell type encoding social interactions. Interestingly, we found that ensembles encoding social interaction and sucrose consumption were distinct, indicating orthogonal representations for social and non-social rewards. To probe the long-term representations of social interactions in NAc, we performed a six-day social interaction assay and found that single-cell NAc social representations were highly dynamic, with only around 25% of neurons similarly encoding social interactions on any two consecutive days. Yet, cross-day decoding analysis showed that neural population representations for social interaction across days were largely stable. Highlighting a role for NAc population stability in controlling social behavior, Cntnap2 NAc short- and long-term population representations were degraded compared to WT controls. Optogenetic inhibition studies showed that D1 and D2-MSNs in NAc played a reciprocal role in controlling social interactions. Importantly, inhibition of NAc improved social interactions in the Cntnap2 model, suggesting a potential therapeutic modality.

We observed large-scale representational drift (*24*, *47–49*), with instability in single neuron activity patterns which nevertheless was associated with stable NAc social representation at population level. In our study, we found the following: 1. There was decreasing overlap of social cells across days; 2. NAc activity on earlier days predicted behaviors on later days; 3. NAc activity on later days predicted behaviors on earlier days. These findings indicate that although NAc neurons dynamically changed their social identities (SE, SI, SN), the whole population consistently carried social information and represented social interactions. Furthermore, the stability of social representations was decreased in Cntnap2^-/-^mice along with reduction of social interaction time as imaging studies showed degraded cross-session decoding performance.

About 95% of neurons in NAc are MSNs that express either D1 dopamine receptors or D2 dopamine receptors (*50*). Canonically, D1-MSNs and D2-MSNs are located in the striatal direct and indirect pathways, respectively. Earlier studies showed that the two types of MSNs play distinct roles in different reward-related behaviors. For example, NAc D1- and D2-MSNs have opposing effects on cocaine reward-seeking behaviors; activation of D1-MSNs promoted cocaine reward-seeking while activation of D2-MSNs suppressed it (*6*). The results of this study are consistent with the idea that D1-MSNs may mediate reinforcement behaviors while D2-MSNs may represent punishment or aversive input (*7*). Our work is consistent with previous study showing that optogenetic stimulation of firing in NAc D1-MSNs promotes social interactions (*10*). Here, our *in vivo* calcium imaging data showed that a larger propotion of D1-MSNs encoding social interaction and D1-MSNs neural activity pattern could better predict social epochs compared to D2-MSNs. In fact, D2-MSNs also play an important role during social interaction. Inhibiting D2-MSNs activity significantly increased interaction time while inhibiting D1-MSNs activity decreased interaction time. Unexpectedly, we found that non-specific inhibition of NAc core neurons increased social interaction time. This is an intriguing finding, as D2-MSNs seem to play an outsized role in controlling social interactions in the NAc, though these neurons are not consistently activated by social interaction. It is possible that these neurons play a strong modulatory role, decreasing social interactions in situations when interactions may be harmful to the animal. The opposing function of D1- and D2-MSNs in social interaction may result from distinct local or downstream circuits activated by each cell type. In addition, dopaminergic (DA) activation may have distinct effects in each cell type. Previous studies found that dopamine acting on D2 receptors potently inhibited NAc neurons while dopamine acting on D1 receptors potentiated glutamatergic drive (*51–53*). Locally applied D2 antagonists increased NAc neuron firing while D1 antagonists decreased cell excitability (*51*). Coactivation of D1 and D2 receptors on NAc neurons causes a reduction in their membrane excitability (*54*). In our case, inhibition of D2-MSNs may block the inhibitory effect caused by DA released in NAc during social interaction which may result in disinhibition of NAc neurons and increased sociability. Moreover, the distinct efferent connectivity of D1- and D2-MSNs may impact how they regulate downstream target, either directly or through feedback connections (*53*, *55*, *56*).

Inhibition of NAc neurons increased social interaction in the Cntnap2^-/-^ model of autism. The effect of inhibition is potentially long-lasting, as we saw increased interaction time during light-OFF sessions that followed light-ON sessions by 96 hours (Fig. 7Q). Therefore optogenetic inhibtion may be causing long-term plasticity in the circuit, which would be advantageous if this strategy is to be adapted for long-term modulation of social behavior in autism.

## Acknowledgments

We thank Z. Donaldson, W. Hong, L. DeNardo for valuable comments on the manuscript and T. Shuman, C. Lee for technical support on miniscope imaging. We also thank R. Hu for project discussion and K. Maguire, C. Yang, Z. Day for experimental assistance.

## Funding

This work was supported by NSF 1707408, U01 NS122124, U01 NS126050. DA-005010, P50HD103557.

## Author contributions

P.Z. and P.G. designed the study. P.Z. performed all experiments and data analysis. X.C. participated in imaging experiments and data analysis. D.A. developed UCLA Miniscope. A.B. set up sucrose consumption equipment and provided code. A.M. performed animal tracking analysis. J.Q. performed post-operative care, behavior test and perfusion. P.Z. and P.G. wrote the manuscript. P.G. supervised the entire project.

## Competing interests

The authors declare no competing interests.

## Data and materials availability

The data and code relating to the paper are available from the corresponding author upon request.

## Notes

### Competing Interest Statement

The authors have declared no competing interest.

