## Supplementary materials for "Accelerated social representational drift in the nucleus accumbens in a model of autism"

**Materials and Methods**

**Mice**

All experimental protocols were approved by the Chancellor's Animal Research Committee of the University of California, Los Angeles, in accordance with the NIH guidelines. All subject mice were male. C57BL6/J mice were obtained from Jackson Laboratory (strain 000664). Cntnap2 mice were obtained from Elior Peles' lab. All mice were maintained on a 12h:12h light/dark cycle with food and water ad libitum (except the mice doing sucrose consumption test). D1-Cre (RRID: MMRRC_017264-UCD) and A2A-Cre (RRID: MMRRC_031168-UCD) mice were donated by Carlos Cepeda's lab at UCLA. These mice are on the C57BL6/J background. Mice were single housed for three to four weeks before *in vivo* calcium imaging and behavior experiments.

**Stereotaxic surgeries**

Mice were anaesthetized with 1 to 2% isoflurane-oxygen mixture and placed into a stereotactic frame (David Kopf Instruments).

For miniscope calcium imaging, 300-500 nl AAV1.Syn.GCaMP6f.WPRE.SV40 was unilaterally injected into NAc core, mPFC (prelimbic cortex, PL), or dCA1 of six- to seven-week-old male C57/BL6J mice at 60 nl $min^{-1}$ (NAc: AP +1.4mm, ML 0.87mm, DV -4.5mm; mPFC: AP +1.8mm, ML 0.4mm, DV -2mm; dCA1: AP -2.1mm, ML 2mm, DV -1.65mm) using a Nanoject microinjector (Drummond Scientific). Five to seven days after virus injection, mice were implanted with a 0.6mm diameter, 7.3mm length relay lens (Inscopix) over NAc, or a 1mm diameter, 4mm length relay lens (Inscopix) over mPFC, or a 1.8mm diameter, 4.7mm length GRIN lens (Edmund Optics) over dCA1 (Coordinates of the center of lens: NAc: AP +1.4mm, ML, 0.9mm, DV -4.3mm; mPFC: AP +1.8mm, ML 0.5mm, DV -1.8mm; dCA1: AP -2.1mm, ML 1.8mm, DV -1.25mm). Lenses were secured to the skull using cyanoacrylate glue and dental cement and covered with Kwik-Sil (WPI). Mice were then individually housed. Two to three weeks later, a miniaturized microscope (UCLA Miniscope, V3) was attached to an aluminum baseplate and placed on top of the GRIN lens. After searching the field of view and adjusting the focal plane, we secured the baseplate with dental cement. A plastic cap was used to cover baseplate.

For the mPFC projection-specific neuron calcium imaging experiment, 500 nl retrograde AAV-SL1-Syn-Cre was injected into downstream brain regions of mPFC (NAc, BLA, MDT or VTA). 500 nl AAV1.Syn.Flex.GCaMP6f.WPRE.SV40 was injected into mPFC. Five to seven days after virus injection, the lens was implanted above the prelimbic cortex. Baseplate implantation was conducted three to four weeks later.

For D1- and D2-MSNs cell type specific calcium imaging, 500 nl AAV1-Syn-Flex-GCamp6f-WPRE-SV40 was injected into the NAc core region in D1- or A2A-Cre mice. Lens implanting and baseplate implantation were the same as mentioned above.

For optogenetic manipulations, 400 nl pAAV-hSyn-EGFP ($2x10^{13}$GC/ml, Addgene 50465) or pAAV-hSyn-eNpHR3.0-EYFP ($2.3x10^{13}$GC/ml, Addgene 26972), pAAV-Syn-DIO-mCherry (1:10 diluted) ($8.4x10^{12}$GC/ml, Addgene 50459) or pAAV-hSyn-SIO-stGtACR2-FusionRed (1:10 diluted) ($1.9x10^{13}$GC/ml, Addgene 105677) was bilaterally injected into NAc core (AP +1.4mm, ML, ±0.87mm, DV -4.5mm) and fiber optical cannulae (Inper) were implanted with 4° tilt angle (AP +1.4mm, ML ±1.3mm, V -4.25mm) and secured by cyanoacrylate glue and dental cement. Photostimulation (continuous light) started four weeks after surgery with four days of habituation before the start.

The locations of viral injections and lens/fiber implantations were confirmed by histological reconstructions. All mice were perfused using 4% PFA. Brains were maintained in PFA for 24-48 hours and transferred into 30% sucrose solution for dehydration. Brains were later sliced to 60 μm-thick slices (Leica cryostat 1950). Brain slices were stained by DAPI and imaged by confocal microscope (ZEISS LSM800 or LeicaDM6 B).

**Behavioral assays**

**Social interaction in the open arena**

Imaged mice were 11-12 weeks old by the time we started the behavioral assay and calcium imaging. All mice for miniscope calcium imaging were habituated to the environment and to wearing a miniscope for three to five days before testing. On the day of the test, mice were allowed to habituate for 30-60 minutes in their home cage inside the experimental room. During the behavioral test, a miniaturized microscope was attached to the baseplate and the mouse was placed into an open arena (45cm X 45cm X 30cm). After the imaged mouse explored the environment alone for one minute, a novel C57BL/6 male mouse, two to three weeks younger than the imaged mouse, was introduced into the arena as a social target. The two mice interacted freely for seven minutes. The imaged mouse was then placed back into its homecage alone for five minutes, after which it was transferred back to the same arena for one minute alone. Then, a 3D printed object (10cm x 10cm x 10cm cube or 10cm diameter, 10cm high cone) was placed in the center of the arena as an object target which the imaged mouse explored for seven minutes. Social interaction sessions and object exploration sessions were counterbalanced randomly. Calcium imaging was performed simultaneously with behavioral recordings performed with a Logitech webcam at 30 Hz.

Mice for conducting optogenetic manipulation during social interaction and object exploration were habituated to the behavior room environment, handling and wearing fibers for four days before tests. During session 1, we tested social interaction under photostimulation (470nm for EGFP and NpHR, 595nm for mCherry and stGtACR2). 72 hours later, we tested object exploration with light OFF as session 2. 24 hours later, we tested social interaction with a novel social target with light OFF as session 3. Another 24 hours later, we tested object exploration with light ON as session 4. During each session, the subject mouse was in the arena for 1-2 mins alone and social interaction/object exploration period lasted 5-9 mins. Behavior was recorded by miniCam (OpenEphys) at 30 Hz.

**Social interaction across sessions**

The imaged mouse was placed into the open arena alone for one minute and then a social target was introduced to it. The same social target was presented during all six days, but the social target was novel to the imaged mouse on the first day. Social interaction lasted for nine minutes. The mouse's behavior was recorded simultaneously with calcium imaging using a WebCam

(Logitech) at a 30 Hz frame rate. To better track the same neurons across days, each mouse was assigned a dedicated miniaturized microscope. The focal plane was adjusted on Day 1 by setting the position of the CMOS sensor and fixing it in place with a set screw.

**Social interaction vs sucrose solution consumption assay**

Mice were water-restricted and trained to lick for sucrose solution (10%) from a metal tube for two to four days while wearing a plastic replica of a miniaturized microscope. Licking triggered release of 15 μl of sucrose solution. Mouse had free access to the sucrose solution for 10 minutes. After each session, each mouse was given access to 1mL water for an hour. The mouse was then placed into the open arena for another ten minutes of habituation. On the day of imaging, the sucrose solution consumption session was conducted first. Calcium imaging was performed with a miniaturized microscope and licking behavior was recorded with a webcam positioned on the side of the mouse. After the sucrose session, the mouse was provided with 1 mL water to drink for one hour. The social interaction session was then conducted in the open arena where the mouse was imaged alone for one minute and then imaged nine minutes while freely interacting with another mouse. The behavior was recorded by a webcam positioned on top of the arena.

**Analysis of behavioral assays**

**Social interaction**

During social or object interaction sessions, the animal's behavior was recorded with a webcam from the top of the arena at 30 frames per second (fps). All videos were manually annotated frame by frame to identify the onset and offset of each epoch of social interaction (or object exploration). We used the following criteria for identifying interaction epochs: 1. The test mouse must conduct active investigation of the social target (or object); 2. Social interaction was defined as approaching, sniffing, grooming, chasing and mounting the social target; 3. For a valid social interaction (object exploration), the interaction onset was identified as the frame in which the nose of the test mouse pointed to the social (object) target within a distance approximately 1/4-1/3 of its body length. The interaction offset was identified as the frame in which the nose of the test mouse started to leave the social (object) target at a distance about approximately 1/4-1/3 of its body length. These behavioral annotations were organized into a binary vector with '1' representing interaction with social target (object target) and '0' representing non-interaction.

For analysis involving aligning and averaging of response to social or object interactions, to allow a sufficient baseline period, we only included interactions when there was at least 5 seconds separating the onset of current epoch and the offset of previous epoch.

**Sucrose consumption**

During sucrose consumption, each frame of the behavioral videos was examined for the presence or absence of licking behavior. To analyze the overlap of licking behavior with imaging, a binary vector was created by Matlab with '1' standing for licking and '0' standing for non-licking.

**Calcium imaging analysis**

To monitor neuronal activity, we performed calcium imaging by UCLA Miniscope (V3). Data was analyzed using MiniscopeAnalysis package (https://github.com/etterguillaume/MiniscopeAnalysis). Raw imaging data was spatially downsampled by a factor of 2. The NoRMCorre algorithm was then applied to perform classic rigid motion correction (*57*). Constrained non-negative matrix factorization for microendoscopic data (CNMF-E) approach was used to identify and extract the spatial shapes and fluorescent calcium activity of individual cells (*58*). All cells' shapes and $Ca^{2+}$ traces were manually inspected to ensure high data quality. Cells with abnormal contours and $Ca^{2+}$ transients were removed. Denoised and demixed traces were used for further analysis.

**Social cell / object cell identification**

Calcium imaging data and behavior videos were recorded simultaneously and synchronized with each other using timestamps derived from the computer's internal clock. Calcium traces were normalized to the maximum value recorded during the entire recording for each neuron. To identify social interaction responsive cells (social cells) and object exploration responsive cells (object cells), we used ROC (receiver operating characteristic) analysis which measures social/object investigation induced response for each neuron (*14, 15*). The overlap of each neuron's activity with social or object interaction epochs was quantified by an area under ROC curve (auROC) value (range 0-1). These values were compared to those obtained when each trace was randomly circularly shuffled 1000 times. Neurons with auROC values over 97.5 percentile of shuffled auROC values were defined as social excited cells (SE cells) and those with auROC values below 2.5 percentile of shuffled auROC values were defined as social inhibited cells (SI cells). Object cells (Object excited cells (OE cells) or object inhibited cells (OI cells)) were also identified in the same way.

**Sucrose cell identification and overlap between social cells and sucrose cells**

Similar to the method of identifying social/object cells, we applied ROC analysis to get the overlap of each neuron's activity with sucrose licking bouts and quantified them by auROC value. We determined the null distribution by circularly shuffling the data 1000 times and then calculated the percentile of each neuron's auROC compared to the null distribution. We then calculated the percentage of overlap between social cells and sucrose cells over the total number of all active cells/social cells/sucrose cells.

**Velocity cell analysis**

To measure the velocity of the imaged mouse, we used DeepLabCut (*33*) to track the nose of the imaged mouse in behavioral videos. To identify neurons whose activity was modulated by velocity, we performed analysis during one-minute epochs before social interaction or object exploration, when the mouse was alone in the arena. The instantaneous velocity during each frame ($v$ ) was calculated from the coordinates of the mouse's nose for each frame ($X_t$, $Y_t$), using

$$v = \sqrt{(Y_{t+1} - Y_t)^2 + (X_{t+1} - X_t)^2}$$

We smoothed this velocity vector using a one-second-long moving window (30 frames), and then computed the correlation (R) between velocity and calcium trace of each cell. Calcium data were then shuffled 1000 times. Correlation between velocity and shuffled calcium trace of each cell ($R_{s1}$, $R_{s2}$, $R_{s3}$,…$R_{s1000}$) were calculated. Cells with an R larger than 97.5% of $R_s$ were identified as positively correlated velocity cells. Cells with an R smaller than 2.5% of $R_s$ were identified as negatively correlated velocity cells. Velocity correlated neurons included both positively and negatively correlated neurons.

**Tracking neurons across days**

To track the same neurons across multiple sessions of calcium imaging, we applied the CellReg package (*37*) (https://github.com/zivlab/CellReg). The spatial footprints of neurons from each session of recording were entered as inputs to CellReg GUI, which then computed a probabilistic model that the spatial footprints were from the same cell across sessions using centroid distances or spatial correlations of the spatial footprints. The mapping of all registered cells to their indices in each session was obtained from CellReg in a matrix of size N (registered cells) x M (imaging sessions). This matrix was used to track the activity of the same cells across different recording sessions.

**Analysis of neural stability across multiple imaging sessions**

Based on the map of all registered cells, we calculated session-wide overlap by counting the number of overlapping cells between Day 1 and Day X and divided it by the total number of cells on Day 1.

$$\text{Session-wide overlap} = \frac{Day1 \cap DayX}{Day1} \text{x}100\%$$

Using ROC analysis to identify socially modulated neurons in each session, another matrix of size n (registered cells) x m (imaging sessions) was created. All cells were assigned their social identity for each session ('+1', SE; '-1', SI; '0', SN; 'NaN', not active). Then we counted the number of overlapping social cells (SE & SI) cells between Day 1 and Day X and divided it by the total number of social cells on Day 1.

$$\text{Social cell overlap} = \frac{Day1Social \cap DayxSocial}{Day1Social} \text{ x } 100\%$$

For calculating the percentage of overlapping social excited cells (SE) and social inhibited cells (SI) among different numbers of sessions, the total number of SE or SI cells on Day 1 was used as the denominator.

**Behavior decoding with population neural activity**

**Single session social interaction/object exploration prediction**

To measure the neural representation of social behavior in different brain regions and different cell types at the population level, we built a decoder based on partial least squares regression (*34*). We used 4-fold cross-validation to evaluate decoder performance. For each iteration, 75% of the data set was used as a training set to train the decoder and testing was conducted on the remaining 25% of the data set. We measured the performance of the decoder by performing ROC analysis and evaluating auROC values as measures of overlap between predicted and actual data. As a control, we also circularly shuffled interaction/non-interaction epoch randomly 50 times and measured the decoder performance for each shuffle. We obtained the chance level of decoder performance by averaging all auROC values of shuffled data.

**Cross-days social interaction prediction**

For multiple sessions calcium imaging data, we also built decoders based on PLSR. In the matrix, to predict behavior on Day Y (column in matrix) using the data on Day X (row in matrix), we used the whole dataset (overlapped cells between Day X and Y) on Day X as the training set to train the decoder and testd on whole dataset (overlapped cells between Day X and Y) on Day Y. To predict same day behavior (Diagonal in matrix, using Day Z data to predict behavior on Day Z), we did 10-fold cross-validation. Chance level decoder performance was determined by averaging 50 auROC values from the shuffled data (randomly shuffle interaction/non-interaction epochs 50 times). For cross-days decoding mPFC calcium data, we randomly selected 30 overlapped cells between any two days and repeated analysis 20 times. For current day decoding (diagonal), we randomly selected 40 cells and repeated analysis 20 times. Matrix results were quantified by averaging the decoder performance according to the number of interval days between training day and test day.

**Statistics**

All statistical analyses were conducted using GraphPad Prism 9 and MATLAB (R2016b). All plots were presented in mean ± SEM unless otherwise specified. Statistical significance was defined with $P<0.05$. All statistical methods and sample numbers are described in individual figure legends.

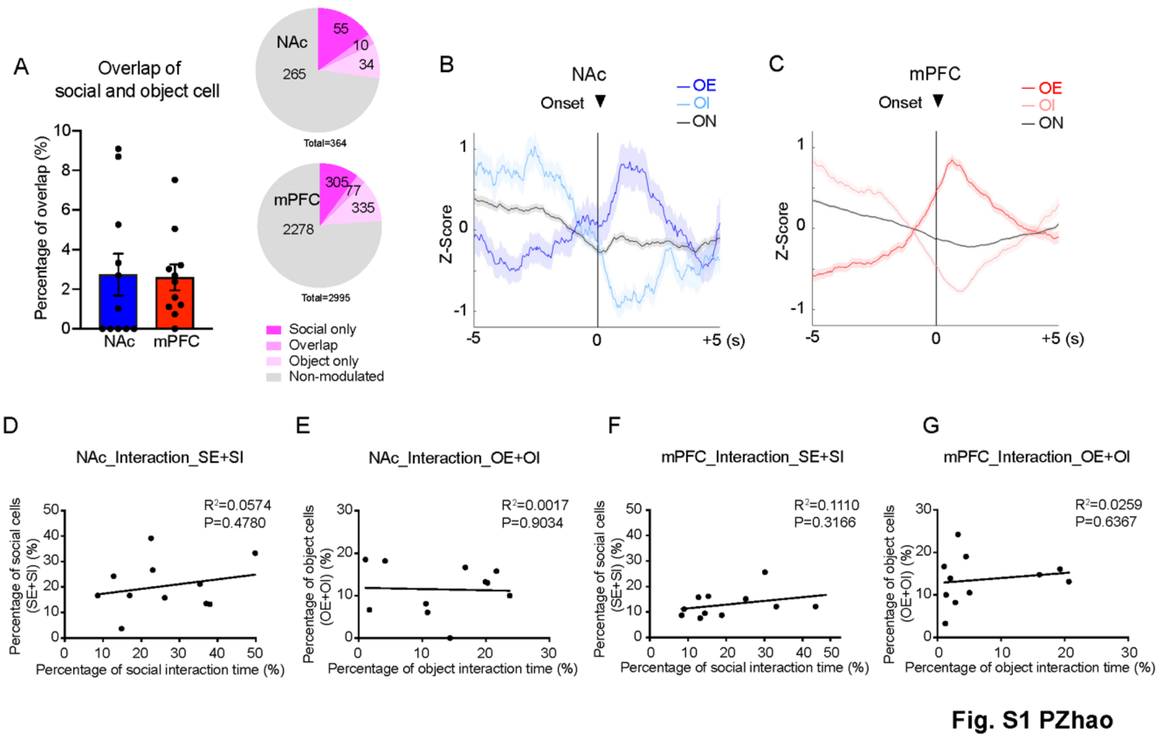


**Fig. S1 Percentage of social interaction and object exploration tuning cells in different brain regions.** (**A**) Overlap of social cells and object cells in NAc and mPFC. NAc, n=11 animals; mPFC, n=11 animals. Mann-Whitney test, P=0.6142. (**B, C**) Average responses of all cells, object excited cells, and object inhibited cells 5 seconds pre- and post-object exploration onset in the NAc and mPFC. (**D, E**) Correlation between the percentage of interaction time and the percentage of social cells (SE+SI) and object cells (OE+OI) in NAc. (**F, G**) Correlation between the percentage of interaction time and the percentage of social cells (SE+SI) and object cells (OE+OI) in mPFC.

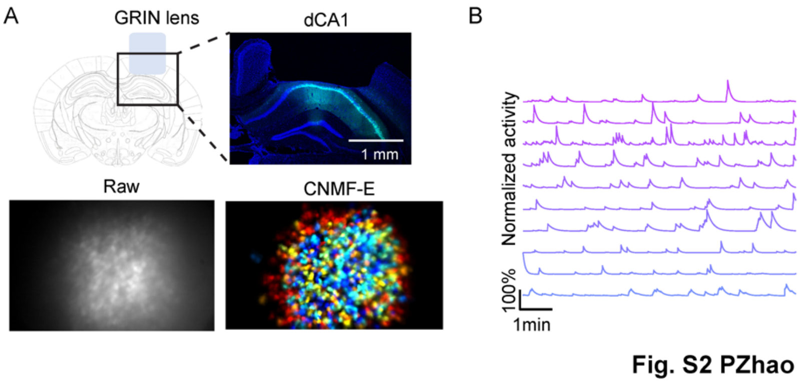


**Fig. S2 Miniscope imaging of dCA1.** (**A**) Schematic of injected virus (Green, GCaMP6f; Blue, DAPI), implanted relay lens for $Ca^{2+}$ imaging in dCA1. Raw image and processed image are shown in lower panel. (**B**) Example of calcium traces extracted from dCA1 imaging.

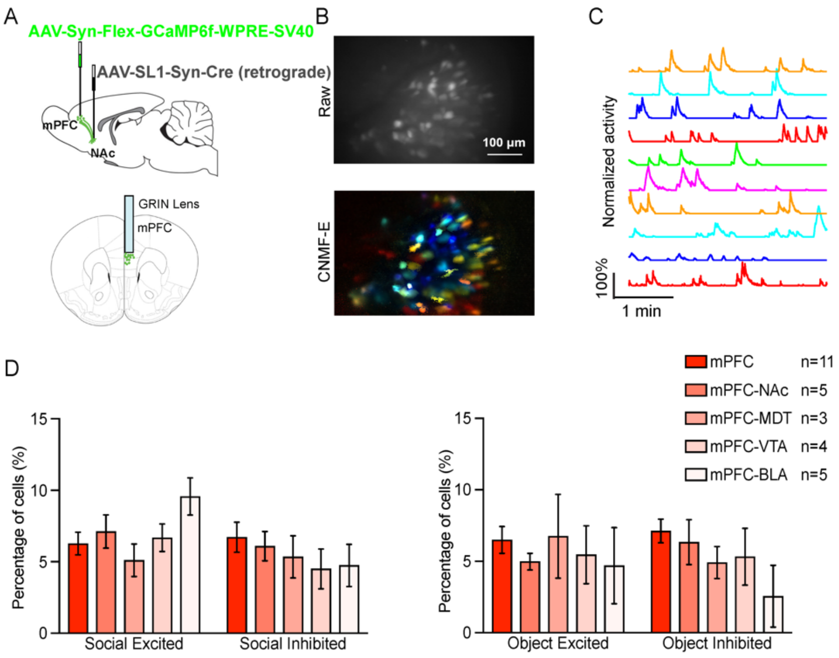


Fig. S3 PZhao

**Fig. S3 Calcium imaging of projection-specific mPFC neurons responding to social interaction and object exploration.** (**A**) Strategy of viral labeling of projection specific mPFC neurons and schematic of lens implantation above mPFC. (**B**) Example of raw calcium fluorescence and extracted single neurons recorded from mPFC-NAc neurons. **(C)** Example of one session calcium traces recorded from mPFC-NAc neurons. (**D**) Summary of percentages of neurons excited/inhibited by social interaction or object exploration in different groups of projection specific mPFC neurons. Two-way ANOVA followed by Tukey's multiple comparisons test. Projection factor, Social, P=0.6546; Object, P=0.2761.

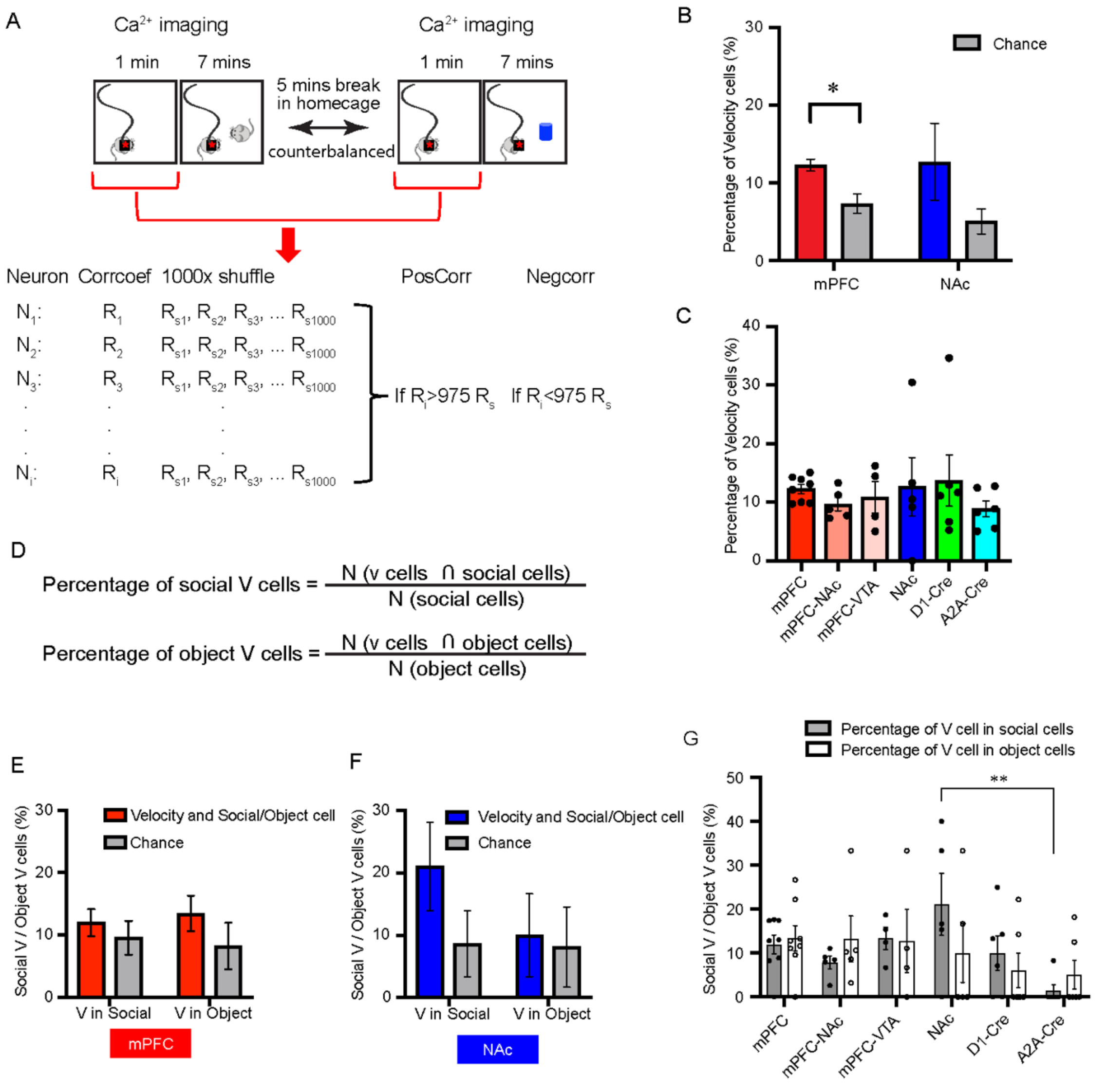


**Fig. S4 PZhao**

**Fig. S4 Velocity cell analysis.** (**A**) Schematic diagram of velocity cell identification. (**B**) Percentage of velocity cells identified in NAc and mPFC compared to chance level. Wilcoxon signed rank test, NAc, n=5 animals, P=0.125; mPFC, n=8 animals, P=0.0156. (**C**) Percentage of velocity cells identified in different groups of cells. One-way ANOVA followed by *Tukey test*. P>0.05 for all. NAc, n=5 animals; mPFC, n=8 animals; D1-Cre, n=6 animals; A2A-Cre, n=6 animals; mPFC-NAc, n=5 animals; mPFC-VTA, n=4 animals. **(D)** Calculating overlap of velocity cells and social/object cells. (**E-G**) Overlap of velocity cells and social/object cells in mPFC and NAc. In E, Two-way ANOVA followed by *Šídák multiple comparisons test.*

Percentage of velocity cells in social cells vs chance, P=0.6473; Percentage of velocity cells in object cells vs chance, P=0.1629. In F, Two-way ANOVA followed by *Šídák multiple comparisons test.* Percentage of velocity cells in social cells vs chance, P=0.4004; Percentage of velocity cells in object cells vs chance, P=0.9763. In G, One-way ANOVA followed by *Tukey test*. NAc vs A2A-Cre (percentage of velocity cells in social cells), *P*=0.0059, all other comparisons, P>0.05. *P<0.05, **P<0.01.

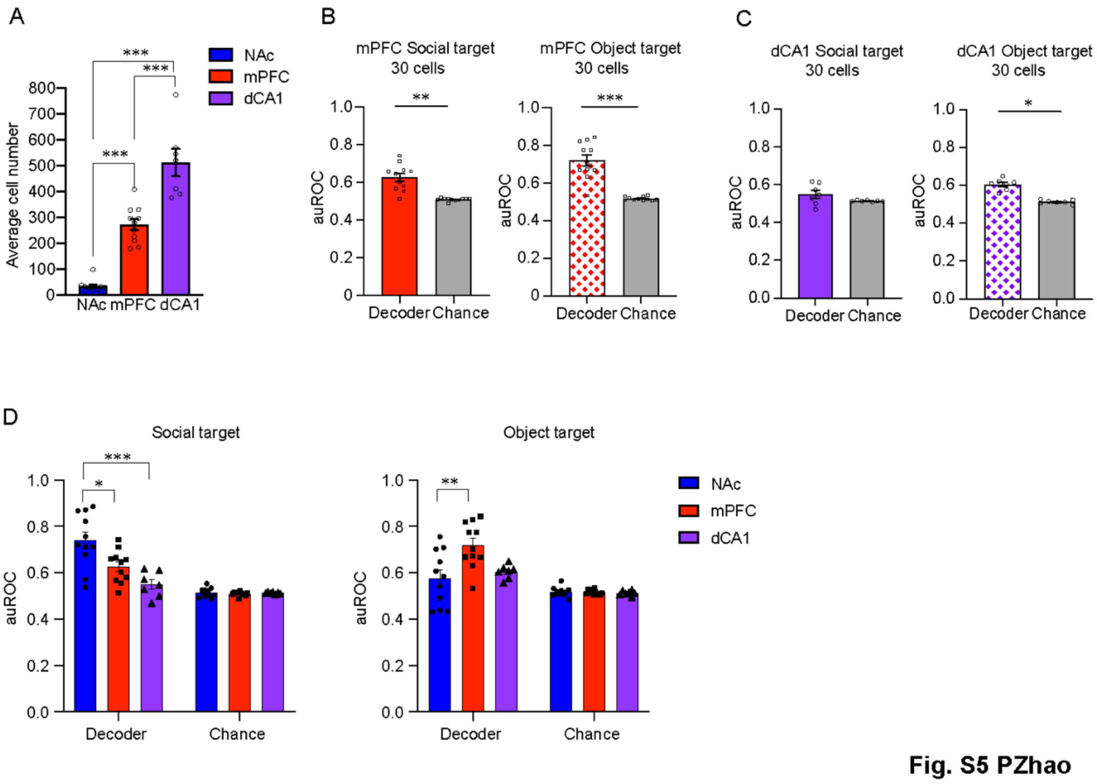


**Fig. S5 Decoder performance of predicting social interaction and object exploration with comparable sizes of training datasets (cell numbers).** (**A**) Average number of cells recoded from NAc, mPFC and dCA1 using Miniscope calcium imaging. NAc, n=11 animals; mPFC, n=11 animals; dCA1, n=7 animals. One-way ANOVA followed by *Tukey test*. P<0.0001 for all. (**B, C**) Decoder performance when predicting social interaction and object exploration with randomly selected 30 mPFC (B) or dCA1 (C) cells (repeated 50 times). Wilcoxon signed rank test. In B, mPFC, social, P=0.002; mPFC, object, P=0.001. In C, dCA1, social, P=0.2188; dCA1, object, P=0.0156. (**D**) Comparison of performance of the decoder trained with comparable numbers of cells' activities in NAc, mPFC, and dCA1. One-way ANOVA followed by *Tukey test*. Social interaction decoding, decoder performance of NAc vs mPFC, P=0.0163, decoder performance of NAc vs dCA1, P=0.0004. Object exploration decoding, decoder performance of NAc vs mPFC, P=0.0047. *P<0.05, **P<0.01, ***P<0.001.

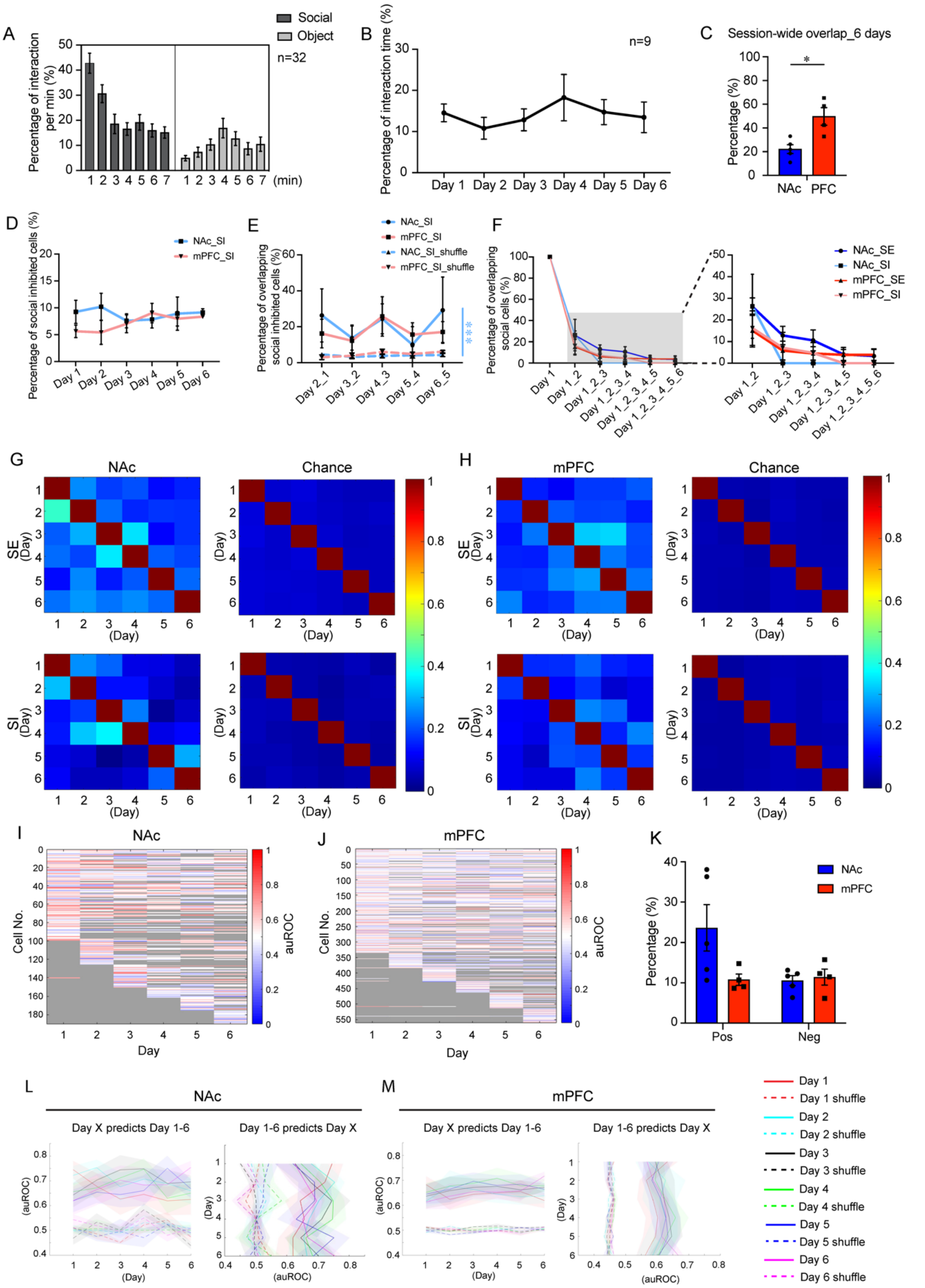


**Science Fig. S6 PZhao**

**Fig. S6 Analysis of social representation across multiple sessions.** (**A**) Adaptation analysis of imaged mice during one session of social interaction/object exploration assay. Percentage of time during each minute spent on investigating social/object target. n=32 animals including NAc, mPFC and dCA1 recorded in animals. (**B**) Percentage of interaction time on each day with the same social target across six days. n=9 animals, including NAc and mPFC recording animals. (**C**) Percentage of overlapped active cells across six days of social interaction in NAc and mPFC. NAc, n=5 animals; mPFC, n=4 animals. Mann-Whitney test, P=0.0317. (**D**) Percentage of social inhibited cells identified in NAc and mPFC on each day. Two-way repeated-measures ANOVA, P=0.4021. (**E**) Percentage of overlapping social inhibited cells between two consecutive days in the NAc and mPFC, compared to chance level. Two-way repeated-measures ANOVA, NAc_SI vs NAc_SI_shuffle, P=0.0002; mPFC_SI vs mPFC_SI_shuffle, P=0.1262. (**F**) Zooming in on the result of Fig. 4H (overlap of social excited/inhibited cells in NAc and mPFC across different length of days). Right panel, Two-way repeated-measures ANOVA, NAc_SE vs mPFC_SE, P=0.3709; NAc_SI vs mPFC_SI, P=0.9415; NAc_SE vs NAc_SI, P=0.1902; mPFC_SE vs mPFC_SI, P=0.2700. (**G, H**) Overlap of social excited/inhibited cells between any two days (denominator is the cell number of the day on the row) in NAc and mPFC. (**I, J**) Single mouse examples of auROC map of NAc and mPFC neurons across six days (grey, cells not present). (**K**) Percentage of cells in NAc and mPFC that consistently positively or negatively respond to social interaction over two days or more. Two-way ANOVA followed by Tukey's multiple comparisons test, Pos NAc vs mPFC, P=0.0877; Neg NAc vs mPFC, P=0.9979; Pos NAc vs Neg NAc, P=0.06; Pos mPFC vs Neg mPFC, =0.9993. (**L, M**) Quantification of decoder performance on crossing days prediction of social interaction in NAc and mPFC according to the result of Fig. 6I, J. Left panel: Decoder performance of using Day X neural activity pattern as training set to predict social interaction on Day 1, Day 2, Day 3, Day 4, Day 5 and Day 6. Right panel: Decoder performance of using neural activities of Day 1, Day 2, Day 3, Day 4, Day 5 and Day 6 as training set to predict social interaction on Day X. *P<0.05, ***P<0.001.

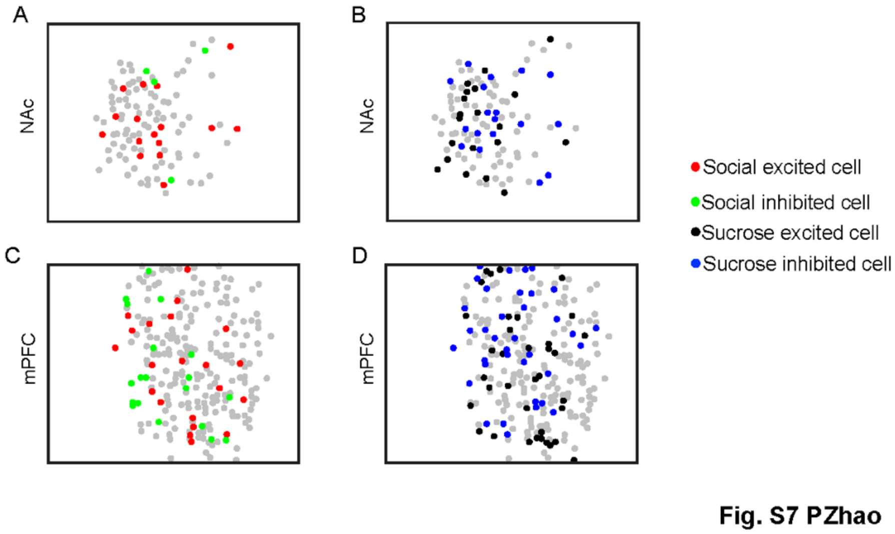


**Fig. S7 Distribution of different tuning cells in NAc and mPFC.** (**A**) Distribution of social excited/inhibited cells in NAc during social interaction. (**B**) Distribution of sucrose excited/inhibited cells in NAc during sucrose consumption. (**C**) Distribution of social excited/inhibited cells in mPFC during social interaction. (**D**) Distribution of sucrose excited/inhibited cells in mPFC during sucrose consumption.

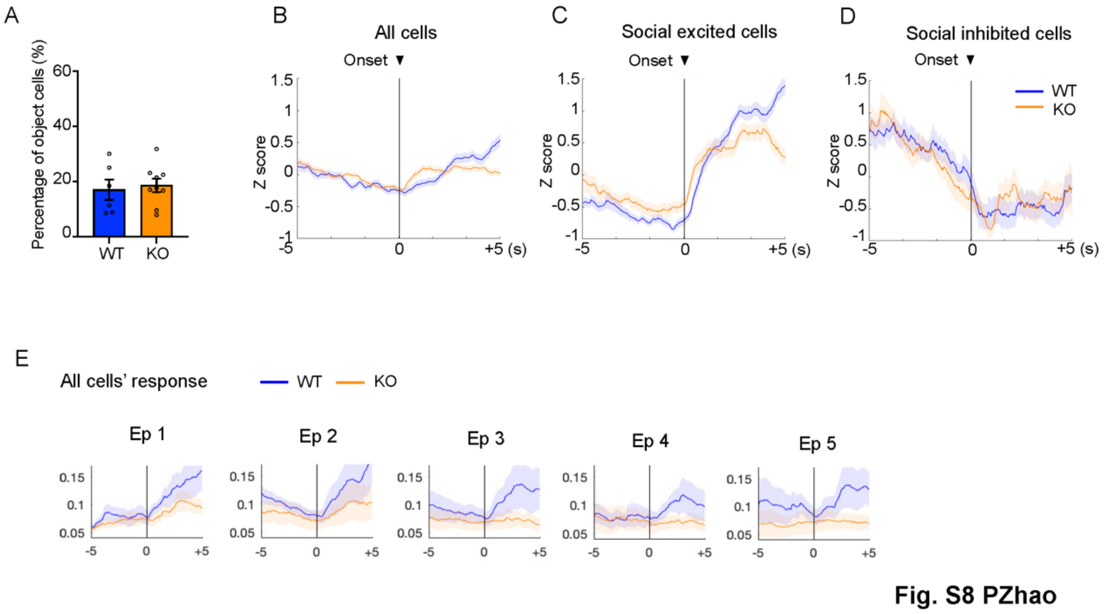


**Fig. S8 Neural activity changes pre- and post-onset of social interaction in Cntnap2$^{-/-}$ mice.** (**A**) Percentage of object exploration modulated cells identified in WT and KO mice in NAc. WT, n=6, animals; KO, n=9 animals. Mann-Whitney test. P=0.8639. (**B**) Averaged activity of all cells recorded from NAc in WT and KO mice aligned by the onset of valid social interaction epochs. (**C**) Averaged activity of social excited cells recorded from NAc in WT and KO mice aligned to the onset of valid social interaction epochs. (**D**) Averaged activity of social inhibited cells recorded from NAc in WT and KO mice aligned by the onset of valid social interaction epochs. (**E**) Average response of all recorded NAc cells in WT and KO mice to the first 5 epochs of social interaction.

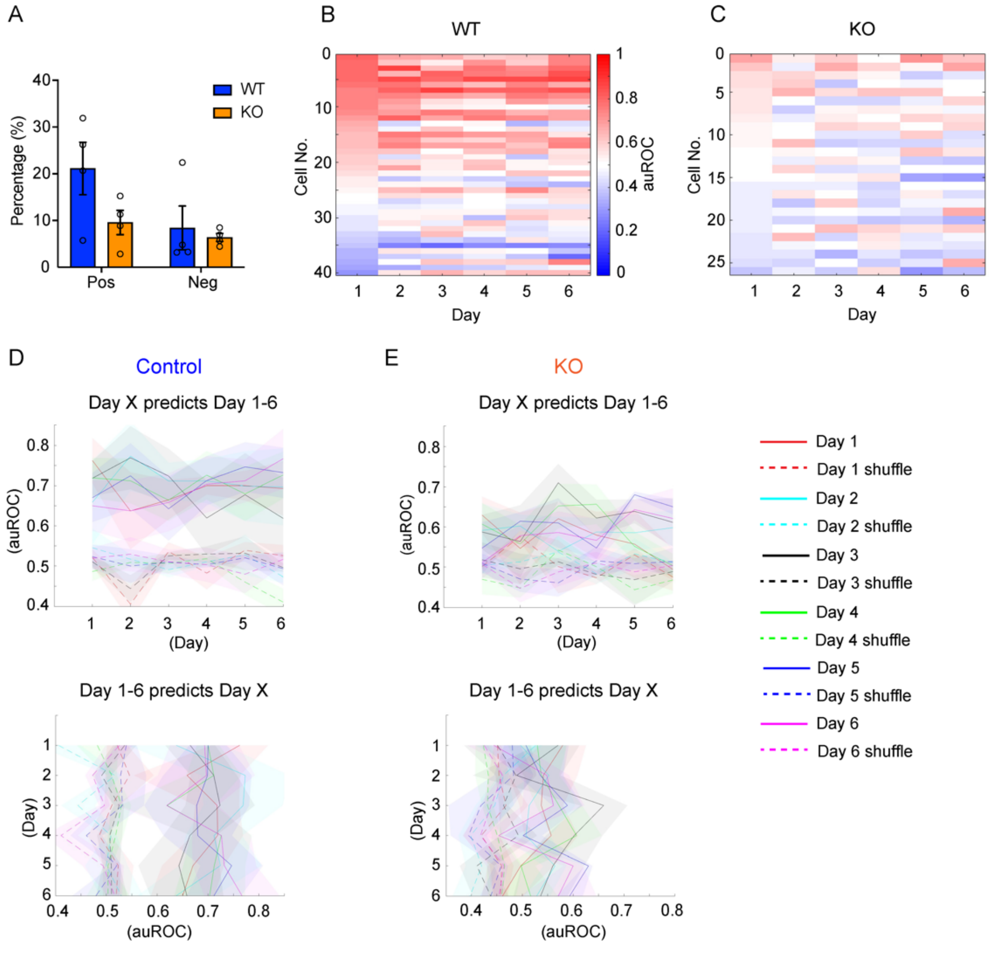


**Fig. S9 Long-term social representation across multiple sessions in Cntnap2$^{-/-}$ mice.** (**A**) Percentage of cells in NAc of WT and KO mice with consistently positive or negative response to social interaction over two days or more. WT, n=4 animals; KO, n=4 animals. Two-way ANOVA followed by Tukey's multiple comparisons test, Pos WT vs KO, P=0.2097; Neg WT vs KO, P=0.9822; Pos WT vs Neg WT, P=0.1520; Pos KO vs Neg KO, P=0.9374. (**B, C**) auROC map of cells present across all six days sorted by Day 1 auROC value. (**D, E**) Quantification of decoder performance on cross-days prediction of social interaction in NAc of control and CNTNAP2 KO mice. Left panel: Decoder performance using Day X neural activity pattern as training set to predict social interaction on Day 1, Day 2, Day 3, Day 4, Day 5 and Day 6. Right panel: Decoder performance using neural activities of Day 1, Day 2, Day 3, Day 4, Day 5 and Day 6 as training set to predict social interaction on Day X.

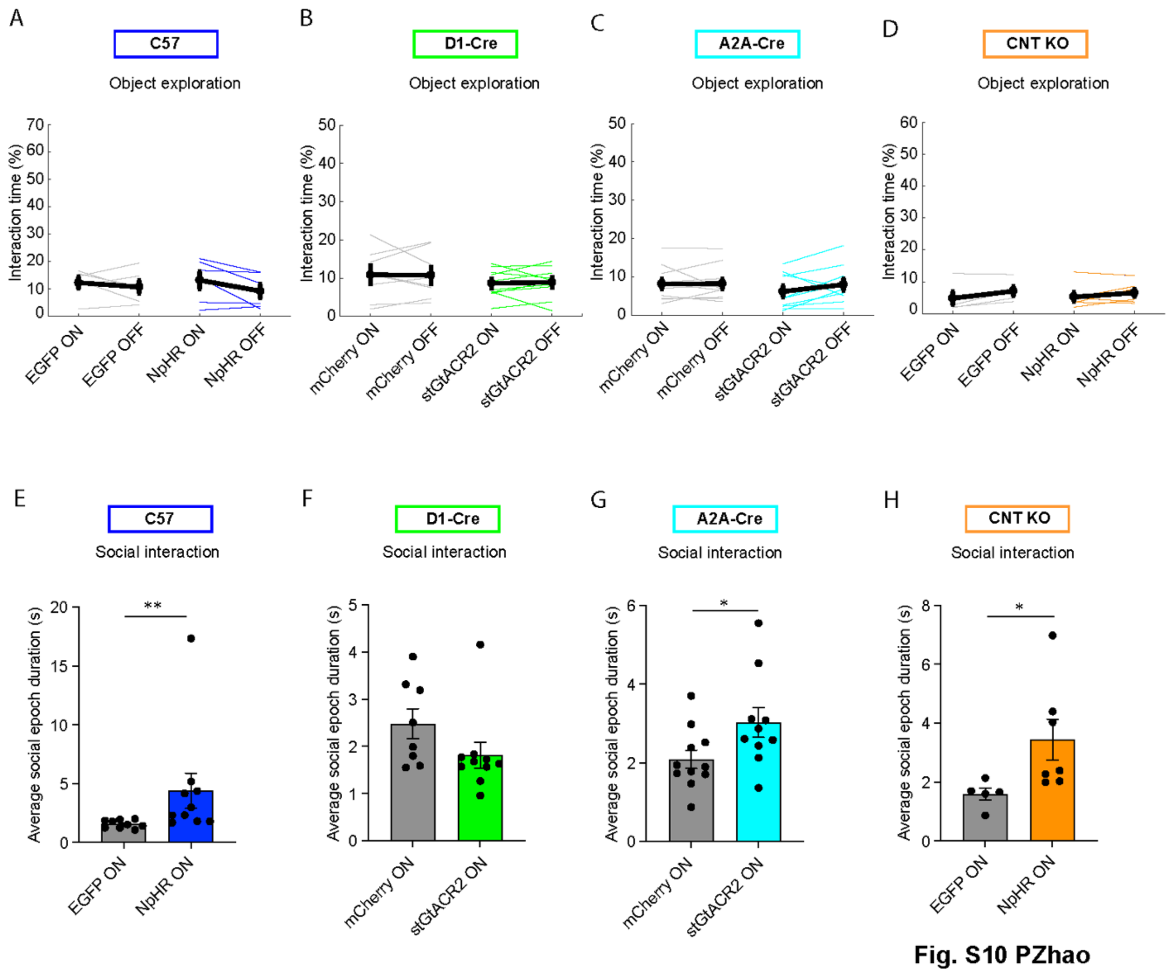


**Fig. S10 Optogenetic manipulation of different cell types in NAc core.** (**A-D**) Optogenetic inhibition of all NAc core neurons in C57 mice (A), D1-MSNs (B), D2-MSNs (C) and all NAc core neurons in Cntnap2$^{-/-}$ mice (D) did not affect object exploration. (**E-H**) Average social interaction epoch duration. Mann-Whitney test, C57 mice, EGFP light ON vs NpHR light ON, P=0.003; D1-Cre mice, mCherry light ON vs stGtACR2 light ON, P=0.1011; A2A-Cre mice, mCherry light ON vs stGtACR2 light ON, P=0.0357; Cntnap2$^{-/-}$ mice, EGFP light ON vs NpHR light ON, P=0.0101. *P<0.05, **P<0.01.
